# Quantifying higher-order epistasis: beware the chimera

**DOI:** 10.1101/2024.07.17.603976

**Authors:** Uthsav Chitra, Brian J. Arnold, Benjamin J. Raphael

## Abstract

Epistasis, or interactions in which alleles at one locus modify the fitness effects of alleles at other loci, plays a fundamental role in genetics, protein evolution, and many other areas of biology. Epistasis is typically quantified by computing the deviation from the expected fitness under an additive or multiplicative model using one of several formulae. However, these formulae are not all equivalent. Importantly, one widely used formula – which we call the *chimeric* formula – measures deviations from a *multiplicative* fitness model on an *additive* scale, thus mixing two measurement scales. We show that for pairwise interactions, the chimeric formula yields a different magnitude, but the same sign (synergistic vs. antagonistic) of epistasis compared to the multiplicative formula that measures both fitness and deviations on a multiplicative scale. However, for higher-order interactions, we show that the chimeric formula can have both different magnitude *and* sign compared to the multiplicative formula — thus confusing negative epistatic interactions with positive interactions, and vice versa. We resolve these inconsistencies by deriving fundamental connections between the different epistasis formulae and the parameters of the *multivariate Bernoulli distribution*. Our results demonstrate that the additive and multiplicative epistasis formulae are more mathematically sound than the chimeric formula. Moreover, we demonstrate that the mathematical issues with the chimeric epistasis formula lead to markedly different biological interpretations of real data. Analyzing multi-gene knockout data in yeast, multi-way drug interactions in *E. coli*, and deep mutational scanning (DMS) of several proteins, we find that 10 − 60% of higher-order interactions have a change in sign with the multiplicative or additive epistasis formula. These sign changes result in qualitatively different findings on functional divergence in the yeast genome, synergistic vs. antagonistic drug interactions, and and epistasis between protein mutations. In particular, in the yeast data, the more appropriate multiplicative formula identifies nearly 500 additional negative three-way interactions, thus extending the trigenic interaction network by 25%.

## 1 Introduction

A key problem in biology is deriving the map from genotype to fitness, or the average reproductive success of a genotype. This map is often referred to as the *fitness landscape* [1]. In the simplest fitness landscape, the fitness of alleles at one locus are independent of the fitness of alleles at every other loci, making fitness either an additive or multiplicative function of the allele at each genetic locus. However, the fitness landscape is complicated by the presence of *epistasis*, or genetic interactions where alleles at one locus modify the fitness effects of alleles at other loci. Epistatic interactions reveal potential functional relationships between genes, as the sign of the interaction (positive or negative) may indicate genetic suppression or functional similarity [2]. The accurate measurement of epistasis is thus crucial for many biological tasks including understanding how genes are organized into genetic pathways [3, 4], modeling protein function and evolution [5, 6, 7, 8, 9, 10, 11], understanding antibiotic resistance [12, 13, 14, 15], and interpreting genome-wide association studies (GWAS) [3, 16, 17, 18, 19].

Over the past few decades, many studies have aimed to measure epistatic interactions from experimental fitness data (reviewed in [3, 4, 5]). Most of these studies measure the simplest type of epistasis: *pairwise epistasis*, or interactions between a pair of genetic loci. Pairwise epistasis is computed by comparing the observed fitness of the double-mutant to the expected fitness under a null model with no epistasis. Almost all formulae for pairwise epistasis use either an *additive* null model, where the expected fitness is the sum *f*_01_ + *f*_10_ of the fitness values of the single-mutants, or a *multiplicative* null model, where the expected fitness is the product *f*_01_*f*_10_ of the fitness values of the single mutants. Under an additive null model, epistasis *ϵ* is typically computed as the difference *ϵ* = *f*_11_− (*f*_10_ + *f*_01_) between observed and expected double-mutant fitness.

For the multiplicative null model, there is notably no agreement in the literature about how to quantify deviation from the null model. In the statistics literature, it is standard to compute multiplicative interaction effects using a *ratio* 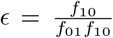 between the observed and expected values (e.g. [20, 21, 22]). On the other hand, many studies in the genetics literature compute epistasis as the *difference ϵ* = *f*_11_− *f*_01_*f*_10_ between observed and expected fitness values of a double-mutant (e.g. [23, 24, 25, 26, 27, 28, 29, 4, 30, 31, 32, 33, 34]). We call the first formula the *multiplicative formula*, as it preserves the multiplicative measurement scale, while we call the second formula the *chimeric formula*, as it measures deviations from a multiplicative model on an additive scale and thus is a “chimera” of additive and multiplicative scales.

The differences between the two pairwise epistasis formulae are not widely appreciated in biological applications. Importantly, we show that the chimeric and multiplicative formula result in different measures of pairwise epistasis, which affects qualitative findings on the strength of an epistatic interaction. At the same time, we also show that the two formulae always yield the same *sign* (or *direction*) of a pairwise interaction. The sign of an epistatic interaction is often the quantity of interest in genetics studies, e.g. negative epistatic interactions are used to quantify functional redundancy [2, 35, 36] and recombination [37, 38, 39, 40]. Thus, the focus of existing literature on the sign of interactions, as well as the focus on pairwise epistasis, may explain why the differences between the multiplicative and chimeric formula are not broadly recognized.

The discrepancies between the multiplicative and chimeric formula are more pronounced and consequential for *higher-order* interactions between three or more loci, which are becoming more widely studied with larger genetic datasets and high-throughput measurements of fitness [41, 2, 42, 43, 44, 45, 46]. Recent studies in yeast genetics [2, 42] and antibiotic resistance [43] independently derived analogous chimeric formula to quantify epistasis between three or more loci and higher-order interactions between antibiotics, respectively, under a multiplicative fitness model. These chimeric formulae were seemingly derived *de novo* and without consideration of the two distinct formula — chimeric and multiplicative — for pairwise epistasis, nor the consequences of conflating multiplicative and additive scales. However, unlike in the pairwise setting, we show that for three or more loci, the chimeric formula is *not* guaranteed to produce the same *sign* of an interaction as the multiplicative formula. Thus, the chimeric formula may indicate a *positive* epistatic interaction while the multiplicative formula shows a *negative* epistatic interaction, and vice-versa. Such inconsistencies raise questions about reported higher-order epistasis in biological applications.

We resolve the mathematical and biological inconsistencies between the different epistasis formulae by deriving a fundamental connection between epistasis and the parameters of the multivariate Bernoulli distribution (MVB), a probability distribution on binary random variables [47]. We demonstrate that this connection to the multivariate Bernoulli is implicit in several other approaches for quantifying epistasis, including the regression models commonly used in GWAS and eQTL analyses [3, 4, 48] and the Walsh coefficients for measuring “background-averaged” epistasis [41, 49, 50]. To our knowledge, the connections we derive between the MVB and the various epistasis formulae have not been previously described.

We leverage the connections to the multivariate Bernoulli distribution to analyze the higher-order chimeric epistasis formulae derived by Kuzmin et al. [2, 42] and Tekin et al. [43]. We show that both the chimeric formulae for pairwise epistasis and the chimeric formulae for higher-order epistasis correspond to the *joint cumulants* of the MVB, a concept from probability theory for measuring interactions between continuous variables [51, 52, 53]. The joint cumulants are known to *not* be an appropriate measure of higher-order interactions for binary random variables [54, 55]. Accordingly, we show that the chimeric epistasis formula are *not* appropriate for measuring higher-order epistasis between biallelic mutations. In this way, just like how the hero Bellerophon in the *Iliad* slayed the monstrous chimera, the multivariate Bernoulli distribution allows us to “slay” the chimeric epistasis formula.

We demonstrate that the mathematical issues with the chimeric epistasis formula lead to markedly different biological interpretations of real data. Analyzing multi-gene knockout data in yeast using the more appropriate multiplicative formula changes the sign of 12% of the 7, 957 trigenic interactions that [2, 42] reported using the chimeric formula.

Many of these sign changes are concentrated on negative interactions, which are more functionally informative than positive interactions and are commonly used to measure functional redundancy between genes [35, 36, 56]. In particular, the multiplicative epistasis formula identifies nearly 500 novel negative interactions that are significantly enriched for several measures of functional redundancy, thus extending the trigenic interaction network by 25%.

We further demonstrate that the multiplicative and additive formulae yield markedly different interactions compared to the chimeric formula in two other applications: the identification of higher-order synergistic and antagonistic drug interactions in *Escherichia coli* and the identification of epistatic interactions between protein mutations in deep mutational scanning experiments. We show that the discordance between the different formulae increases with interaction order: the additive formula shows significantly less antagonism between five-way interactions compared to the chimeric formula used in [57], while for some proteins there is up to substantial (up to 60%) disagreement in the sign of interaction between the multiplicative and chimeric formulae.

## 2 Results

### 2.1 Pairwise epistasis: additive, multiplicative, and chimeric

Pairwise epistasis describes interactions between two genetic loci. We assume that each locus is *biallelic*, i.e. each locus has two alleles labeled 0 and 1. Thus for a pair of loci there are four possible genotypes: the wild-type 00, the single mutants 01 and 10, and the double mutant 11. Accordingly, for a pair (*i, j*) of loci there are four corresponding fitness values: the wild-type fitness *f*_∅_, corresponding to the wild-type genotype 00 with no mutations; the single-mutant fitnesses *f*_*i*_, *f*_*j*_, corresponding to the genotypes 01 and 10 with either locus *i* or locus *j* mutated, respectively; and the double-mutant fitness *f*_*ij*_, corresponding to the genotype 11 with both loci mutated. Pairwise epistasis is measured by comparing the observed double-mutant fitness *f*_*ij*_ to the expected fitness under a null model with no epistasis.

In practice, the fitness *f* of a genotype, i.e. the mean reproductive success of the genotype, cannot be directly measured. Instead, experiments typically measure traits that are expected to be highly correlated with fitness, e.g. cellular reproductive or growth rate, and are assumed to follow either an additive or multiplicative scale [58]. Accordingly, the two standard null models of fitness for measuring epistasis are the *additive* model and the *multiplicative* model.

In the *additive* model, mutations are assumed to have an additive effect on fitness [29, 4, 3]. Thus the expected double-mutant fitness is given by *f*_*i*_ + *f*_*j*_, or the sum of fitness effects of the individual mutations, under the usual assumption that fitness values are normalized such that the wild-type fitness *f*_∅_ = 0. The additive model has been used in studies of drug resistance [59, 60, 61, 62, 63], protein binding [50, 41], and quantitative genetics [4, 28, 64]. The pairwise epistasis measure 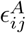 under the additive model is equal to the difference between the observed and expected double-mutant fitness values:

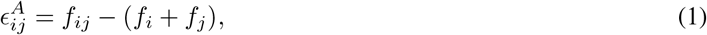

and the sign of the interaction (i.e. positive vs. negative) is given by the sign sgn 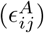 of the epistasis measure 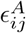.

In the *multiplicative* fitness model, mutations are assumed to have a multiplicative effect on fitness. Thus, the expected double-mutant fitness is given by *f*_*i*_ · *f*_*j*_, or the product of fitness effects of the individual mutations, under the typical assumption that fitness values are normalized such that the wild-type fitness *f*_∅_ is equal to 1. The multiplicative fitness model has been used to model cellular growth rates [4, 23, 33, 2, 31, 65, 66, 67] and protein structure [68]. The pairwise epistasis measure 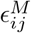 under the multiplicative fitness model is equal to the ratio between the observed and expected double-mutant fitness values:

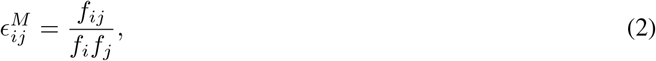

and the sign of the interaction is determined by whether the multiplicative measure 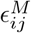 is greater than or less than 1.

The additive and multiplicative fitness models are closely related: if fitness values *f* are multiplicative, then the *log-*fitness values log *f* are additive. Thus, the sign of an interaction under the multiplicative model is also given by the sign sgn(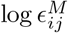) of the log-epistasis measure 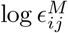. We prove that the additive and multiplicative epistasis measures are closely related to two other notions of epistasis used in the genetics literature: the linear/log-linear regression frameworks [3, 21] and the Walsh coefficients [41, 49, 50, 60, 69]. See Methods for details.

Curiously, there is a third epistasis formula that is widely used for the multiplicative fitness model. Here, deviations from the multiplicative model are measured on an additive scale, resulting in the following *chimeric* formula for pairwise epistasis:

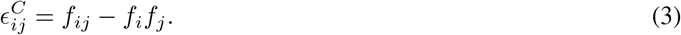

We refer to 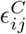 as the chimeric epistasis measure because it measures deviations from a multiplicative null model on an additive scale, and is thus a *chimera* of both the multiplicative and additive measurement scales. Although the chimeric epistasis measure quantifies deviations from the multiplicative model, the sign of the interaction is given by the sign sgn 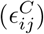of the chimeric measure 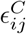 as in the additive fitness model. The chimeric measure 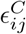 is widely used in the genetics literature (e.g. [23, 24, 25, 26, 27, 28, 29, 4, 30, 31, 32, 33, 34]) and in the drug interaction literature (e.g. [70, 71, 72, 73, 74, 75, 57, 43, 76, 77]).

The chimeric epistasis measure 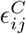 is widely used because of its interpretation as a *residual*, i.e. the difference between the observed and expected values of a measurement. However, despite the simplicity of this explanation, it is not statistically sound, as residuals are only appropriate for additive models. For multiplicative models, instead it is standard to compute the *ratio* between observed and expected measurements, rather than the difference [21]; here, the ratio between observed and expected fitness corresponds to the multiplicative epistasis measure 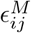. Indeed, Wagner [78, 22] notes that preserving the multiplicative measurement scale (by using the ratio) is required in order to guarantee meaningful notions of statistical and functional interactions.

The differences between the multiplicative epistasis measure 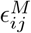 and chimeric epistasis measure 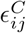 do not appear to be widely appreciated in either the applied or theoretical literature. Almost every biological study that uses the chimeric epistasis measure 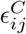 does not consider the multiplicative measure 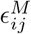. On the other hand, while many in the statistics literature draw a distinction between additive and multiplicative interaction effects (e.g. [21, 22]), none of these papers discuss the chimeric interaction measure 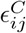 that is frequently used in the genetics and drug interaction literature. An exception is Gao, Granka, and Feldman [79] who generously refer to the multiplicative formula 2 as a *“rescaling of [the chimeric] formula”*. However, we take the stronger view that “rescaling” is a generous term that obscures consequential implications of the two formula.

While both the chimeric measure 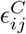 and the multiplicative measure 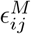 are described as measuring deviations from a multiplicative fitness model, the two measures are not equal. In particular, the (log-) multiplicative epistasis measure 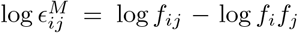 computes the difference between the observed and expected double-mutant fitness values on a logarithmic scale (Figure 1A) while the chimeric epistasis measure 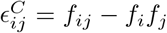 computes the difference directly (Figure 1B). Thus, the chimeric epistasis measure 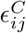 may over- or under-state the strength of a pairwise interaction in a multiplicative fitness model, as we demonstrate numerically (Supplementary Text). We note that when the double-mutant fitness *f*_*ij*_ and single-mutant fitness values *f*_*i*_, *f*_*j*_ are close to 1, the chimeric measure 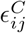 is approximately equal to the log-multiplicative measure 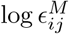(Supplementary Text).

**Figure 1:**
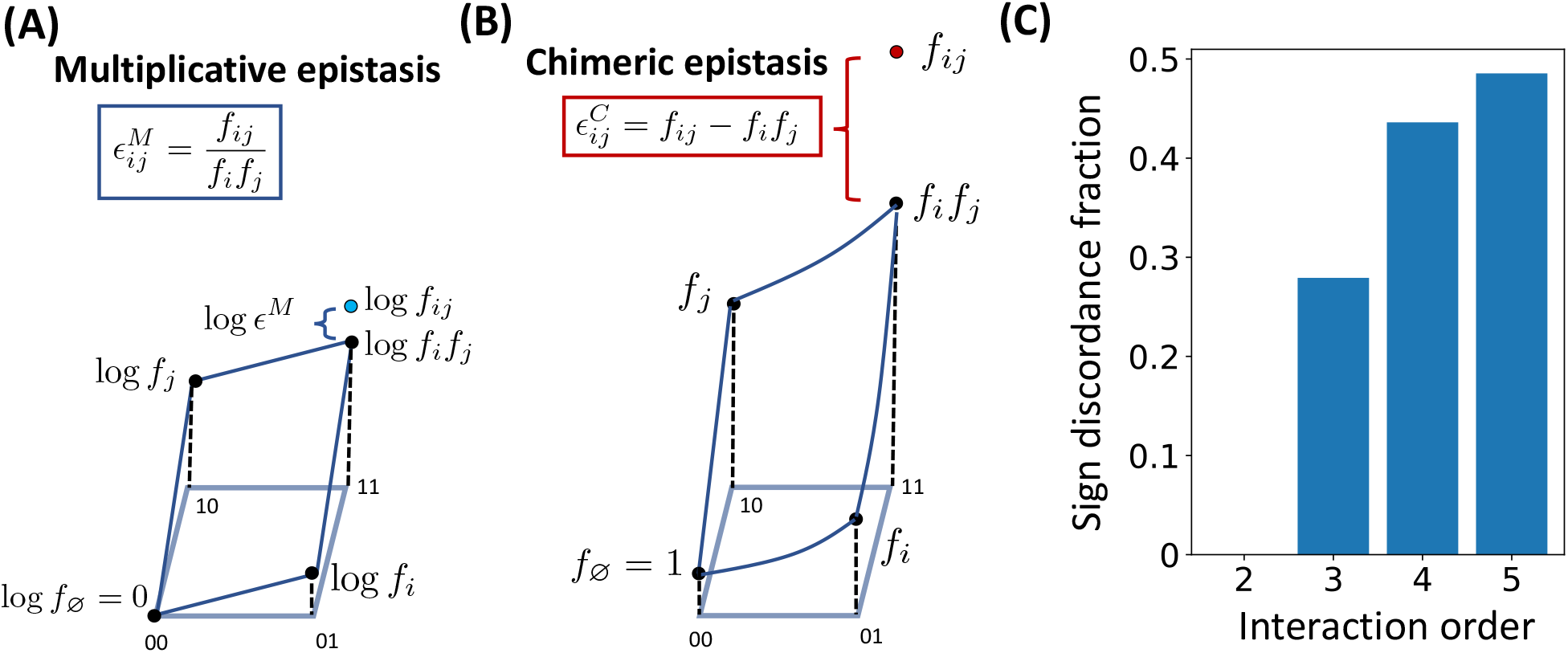
**(A)** For a pair (*i, j*) of loci, the multiplicative epistasis measure 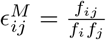 is the *ratio* between the observed fitness *f*_*ij*_ of the double mutant and the expected fitness *f*_*i*_*f*_*j*_ under a multiplicative null model. Equivalently, the logarithm log 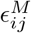 of the epistasis measure is given by 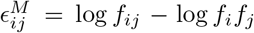, or the difference between the observed and expected values in log-fitness space. **(B)** The chimeric epistasis measure 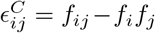 is the *difference* between the observed and expected fitness values of the double-mutant under a multiplicative fitness model. Thus, the chimeric measure 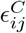 is a different measure of pairwise epistasis that conflates both multiplicative and additive scales. **(C)** The fraction of instances where the signs sgn(log *ϵ*^*M*^) and sgn(*ϵ*^*C*^) of the multiplicative and chimeric fitness formula, respectively, disagree (“sign discordance fraction”) for interaction orders *L* = 2, …, 5, where fitness values *f*_*i*_, *f*_*ij*_, … are sampled uniformly at random from the interval [0, 1]. For two loci, the sign of the two measures always agree (see Proposition 1), but for more than two loci, there is substantial disagreement.

Nevertheless, we prove (Methods) that the chimeric measure 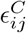 has the same *sign* of an interaction as the multiplicative measure 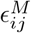, i.e. the chimeric measure and the multiplicative measure agree on the *direction* of deviation from the null model but not the *magnitude*. Thus, using either the chimeric or multiplicative measures will not affect findings that depend on the *sign* of an epistatic interaction, such as the relationship between increased recombination and negative epistasis [37, 40]. However, we emphasize that while in many applications the direction of deviation is the quantity of interest, the magnitude of the deviation (i.e. effect size) is important for statistical tests.

However, the fact that the two measures agree on the sign of an interaction is true only for pairwise epistasis and not higher-order epistasis, as we will show next.

### 2.2 Higher-order epistasis

For higher-order epistasis, or interactions between three or more genetic loci, we find that the difference between the multiplicative and chimeric epistasis measures are much more consequential. Under the multiplicative fitness model, the three-way epistasis measure 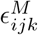 between loci *i, j, k* is given by the ratio between observed and expected triple-mutant fitness:

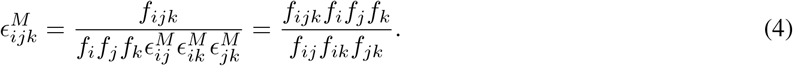

Recent work in the yeast genetics [2, 42] and drug interaction [43] literature claim to use a multiplicative fitness model, but derive a different formula to quantify deviations between the observed and expected fitness for three loci:

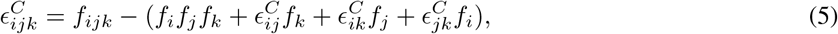

where 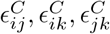 are the pairwise chimeric epistasis measures in (3). Note that as in the pairwise case, formula (5) mixes the additive and multiplicative scales in a complex manner. Thus, we refer to 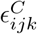 as the *chimeric* three-way epistasis measure.

As in the pairwise setting, the three-way chimeric measure 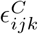 in (5) is clearly different from the three-way multiplicative measure 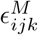 in (4). However, we show (Figure 1C) these formula may differ in both the magnitude of epistasis (as in the pairwise setting) *and* in the *sign* of epistasis. Thus, one formula may indicative positive epistasis between three loci while another formula may indicate negative epistasis, and vice-versa. We demonstrate this numerically (Supplementary Text), showing that even when there is no three-way epistasis according to a multiplicative null model (i.e. 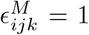), the chimeric three-way epistasis measure 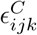 may incorrectly indicate either positive or negative three-way epistasis. Moreover, on simulated data, we find that the difference between the two formula may be quite pronounced with approximately 28% of triples having different signs of epistasis between the two epistasis formula (Figure 1C).

Tekin et al. [43] extended the three-way chimeric epistasis formula (5) to compute a 4-way chimeric epistasis measure 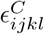 and a 5-way chimeric epistasis measure 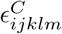. We find even more substantial differences in the sign of epistasis between these 4-way and 5-way chimeric epistasis measures and the 4-way and 5-way multiplicative epistasis measures (equation (18) in Methods). On simulated data, only approximately 57% of the 4-way and 52% of the 5-way interactions have the same sign of higher-order epistasis using both the chimeric and multiplicative epistasis formulae (Figure 1C).

This substantial disagreement between the chimeric and multiplicative epistasis measure motivates a deeper mathematical understanding of the various epistasis formulae, which we undertake in the next section.

### 2.3 Unifying epistasis measurements with the multivariate Bernoulli distribution

A genotype of biallelic mutations on *L* loci can be represented as a binary string of length *L*, where 0 corresponds to the wild-type allele and 1 corresponds to the mutant, or derived, allele. For example, the string 01100 represents the genotype of *L* = 5 loci with mutations in the second and third loci. The fitness values of all genotypes, often referred to as the fitness landscape, corresponds to a function *f* that maps a binary string **x** ∈ {0, 1}^*L*^ to its fitness *f*_x_.

A natural approach for studying a fitness landscape function *f* is to view it as a *distribution* on the set {0, 1}^*L*^ of binary strings, where the probability *p*_x_ of a binary string **x** is proportional to its fitness *f*_x_. Such distributions are often used by protein structure models [80, 81]. Moreover, modeling fitness as a probability has a natural biological interpretation: if the growth rate of an organism with genotype **x** is given by its fitness *f*_x_, and if there are initially an equal number of organisms of each of the 2^*L*^ genotypes **x** ∈ {0, 1}^*L*^, then after one unit of time the frequency *p*_x_ of each genotype **x** will be proportional to its fitness *f*_x_.

Here, we model the fitness landscape using the *multivariate Bernoulli* (MVB) distribution [47, 82] which describes *any* distribution on the set {0, 1}^*L*^ of binary strings. Formally, a multivariate random variable (*X*_1_, …, *X*_*L*_) distributed according to a MVB is parametrized by the probabilities *p*_x_ = *P* ((*X*_1_, …, *X*_*L*_) = **x**) for each binary string **x** = (*x*_1_, …, *x*_*L*_) ∈ {0, 1}^*L*^. We model the genotype (*X*_1_, …, *X*_*L*_) of an organism as a random variable distributed according to a MVB parametrized by the probabilities 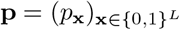.

We prove that the additive, multiplicative, and chimeric measures of epistasis – as well as the Walsh coefficients described in [50, 41, 49, 60, 69] – correspond to different parametrizations of the MVB distribution (Table 1, Methods). We briefly describe these results below.

**Table 1:** Relationship between the different measures of epistasis and the parametrizations of the multivariate BernFoulli distribution.

| | Fitness values $f$ | Parameters of multivariate Bernoulli distribution |
| --- | --- | --- |
| Additive epistasis measure $\epsilon^A$ | Log-probabilities $\log p$ | Natural parameters $\beta$ |
| Multiplicative epistasis measure $\epsilon^M$ | Probabilities $p$ | Natural parameters $\beta$ |
| Chimeric epistasis measure $\epsilon^C$ | Moments $\mu$ | Joint cumulants $\kappa$ |
| Walsh coefficients | Probabilities $p$ | Moments of $(1 - 2X_1, \dots, 1 - 2X_L)$ |

#### Multiplicative and additive epistasis

Suppose the fitness values *f*_x_ ∈ ℝ of each genotype **x** = (*x*_1_, …, *x*_*L*_) ∈ {0, 1}^*L*^ are proportional to the corresponding probability *p*_x_ of a multivariate Bernoulli random variable (*X*_1_, …, *X*_*L*_), i.e. *f*_x_ = *c* · *p*_x_ for some *c >* 0. We prove that the (log-) multiplicative epistasis measures are equal to the *natural parameters* of the MVB. The natural parameters ***β*** = {*β*_*S*_} _*S*⊆{1,…,*L*}_ are another parameterization of the MVB that encode conditional independence relations between the random variables *X*_1_, …, *X*_*L*_; see [47, 83]. We prove a similar result for the additive epistasis measure under the assumption that the fitness *f*_x_ is proportional to the *log-*probability log *p*_x_. See Methods and Supplementary Text for theorem statements and proofs.

Our theoretical results provide a novel connection between the multiplicative epistasis measure and interaction coefficients in a log-linear regression model. This is because for each subset *S* of loci, the natural parameter *β*_*S*_ corresponds to the interaction term for the subset *S* in a log-linear regression model [82, 83, 47]. Such interaction terms are a standard approach for measuring epistasis in genetics, e.g. GWAS or eQTL analyses for quantitative traits [3, 4, 48].

We also prove that the natural parameters ***β*** of the MVB are closely related to the two standard approaches for measuring pairwise SNP-SNP interactions in a case-control GWAS: logistic regression and conditional independence testing [84]. Specifically, we prove that the interaction term in a logistic regression is equal to a 3-way interaction term *β*_*ijk*_ in a MVB, while the conditional independence test is equivalent to testing whether a 2-way interaction term *β*_*ij*_ and a 3-way interaction term *β*_*ijk*_ are both equal to zero. These interaction terms are equal to the corresponding log-multiplicative epistasis measures log *ϵ*^*M*^. To our knowledge, the relationship between the multivariate Bernoulli, logistic regression, and conditional independence testing has not been explicitly described previously in either the genetics or statistics literature.

Thus, our results show that the additive and multiplicative epistasis measures are implicitly computing interaction terms in regression models commonly used in genetics.

#### Chimeric epistasis

The connection between the epistasis formulae and the MVB distribution allows us to derive, to our knowledge, the first mathematically rigorous definition of the chimeric epistasis formula. Specifically, suppose the fitness value *f*_x_ of each genotype **x** = (*x*_1_, …, *x*_*L*_) is equal to a corresponding *moment* 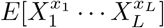 of the random variable (*X*_1_, …, *X*_*L*_). Then we define the chimeric epistatic measure 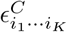 as the *K*-th order *joint cumulant* 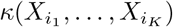 of the random variables 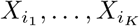(Table 1). Joint cumulants are a concept from probability theory that are used to quantify higher-order interactions between random variables [51, 52, 53]. See Methods for a formal definition.

We emphasize that prior literature on higher-order interactions does not rigorously define the chimeric epistasis measure. For example, Kuzmin et al. [2, 42] does not explicitly state the connection between the joint cumulant and their three-way chimeric formula, while Tekin et al. [43] heuristically uses the joint cumulant formulae without specifying random variables or a probability distribution — thus obscuring any assumptions made by using joint cumulants to measure higher-order interactions.

Our explicit definition of the *K*-th order chimeric epistasis measure 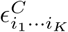 as the *K*-th order joint cumulant reveals two critical issues with the chimeric formula. First, the assumption that the fitness values *f* are equivalent to the moments of a MVB random variable is not biologically reasonable for higher order interactions between three or more loci. This is because the moments assumption implies that the fitness of a particular genotype depends on the probability of many other genotypes. For example, if we assume that the fitness values for *L* = 4 loci are moments of the MVB, then the fitness *f*_1100_ of a double mutant is equal to the moment *E*[*X*_1_*X*_2_], which is equal to

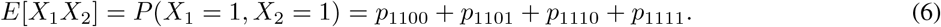

However, it is not clear why the fitness *f*_1100_ of a *single* genotype, 1100, should equal the sum of the probabilities of *four different* genotypes, 1100, 1101, 1110, and 1111.

The second issue is that joint cumulants are not an appropriate measure of higher-order interactions between *binary* random variables. The differences between the joint cumulants and natural parameters ***β*** have been previously investigated in the neuroscience literature, as both quantities have been used to quantify higher-order interactions in neuronal data. For example, Staude et al. [54] write that the joint cumulants *κ* and natural parameters ***β*** *“do not measure the same kind of dependence. While higher-order cumulant correlations indicate additive common components* … *the [natural parameters] directly change the probabilities of certain patterns multiplicatively”*. In particular, the natural parameters ***β*** measure *“to what extent the probability of certain binary patterns can be explained by the probabilities of its sub-patterns”* [54]. Thus, for biallelic genotype data, the natural parameters ***β*** correspond exactly with the epistasis we aim to measure, i.e. how the fitness of a binary pattern can be explained by the fitness of its “sub-patterns”, while the joint cumulants do not.

### 2.4 Simulations

#### 2.4.1 Multiplicative fitness model

We performed simulations to demonstrate the discrepancy between the multiplicative epistasis measure and the chimeric epistasis measure. Since both the multiplicative and chimeric measures use a multiplicative fitness model, we simulated fitness values **f** for *L* = 10 loci following a multiplicative fitness model with *K*-way interactions *β* for different choices of interaction order *K*, and with multiplicative Gaussian noise with standard deviation *σ* (Methods). We computed the *K*-way multiplicative measure 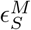 and chimeric measure 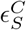 for each set *S* ⊆ {1, …, *L*} of loci of size |*S*| = *K*, and we compared these two measures to the true interaction measure *β*_*S*_.

We first assessed whether the *sign* of the epistasis measures, i.e. sgn 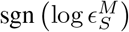and sgn 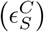, match the sgn(*β*_*S*_) of the true interaction term *β*_*S*_, since the sign of a measure indicates whether there is a positive or negative interaction between mutations in the loci *S*. We observed (Figure 2A) that for pairwise interactions (*K* = 2), both the multiplicative measure *ϵ*^*M*^ and chimeric measure *ϵ*^*C*^ have the same sign as the true interaction measure *β* for the same fraction of instances, which matches our theoretical result (Proposition 1, Methods). However, for higher-order interactions (*K >* 2), the chimeric measure *ϵ*^*C*^ has an incorrect sign more often than the multiplicative measure *ϵ*^*M*^ (Figure 2A). In particular, for *K* = 5-way interactions, even with no noise (i.e. *σ* = 0), the chimeric measure has a different sign than the true interaction parameter *σ* for more than 30% of simulated instances. We also highlight that when there is no noise, i.e. *σ* = 0, the multiplicative measure always has the same sign as the true interaction parameter *β*, i.e. sgn(log *ϵ*^*M*^) = sgn(*β*), which agrees with Theorem 1.

**Figure 2:**
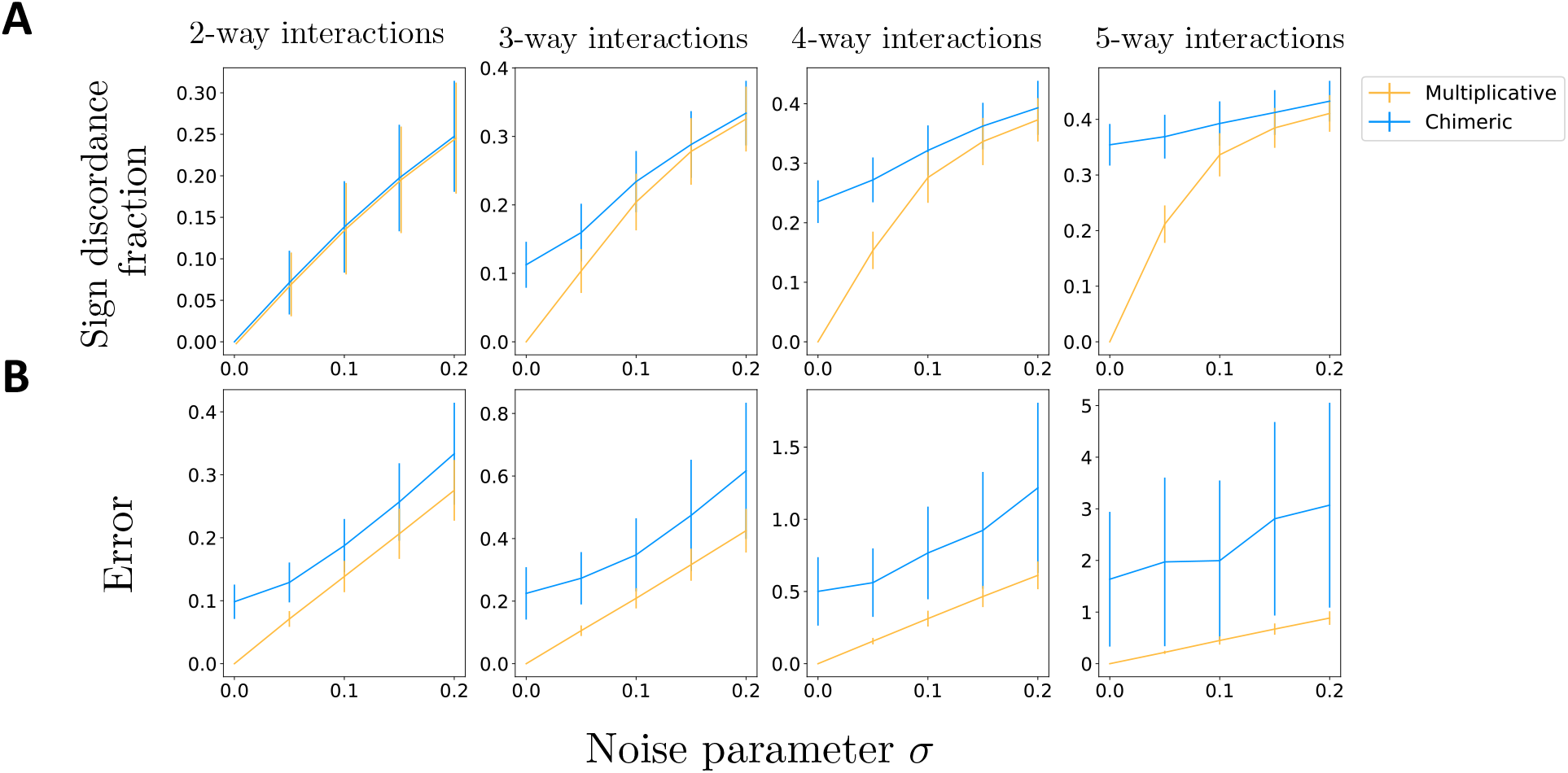
Fitness values **f** are simulated following a multiplicative fitness model with interaction parameters *β*, with maximum interaction order *K* = 2, …, 5, and multiplicative Gaussian noise with standard deviation *σ*. **(A)** The fraction of *K*-way interactions where the sign of the log-multiplicative epistasis measure log *ϵ*^*M*^ (orange) and the chimeric epistasis measure *ϵ*^*C*^ (blue) do not match the sign of the true interaction parameter *β*. **(B)** The average absolute difference (“error”) |*β*−log *ϵ*^*M*^| and |*β*− *ϵ*^*C*^| between the true interaction parameter *β* and (orange) the log-multiplicative measure log *ϵ*^*M*^ and (blue) the chimeric measure *ϵ*^*C*^, respectively. For all panels, these quantities are averaged across 100 simulated fitness values.

We next compared how well the *magnitudes* of the multiplicative and chimeric epistasis measures agree with the magnitude of the true interaction parameters. We computed the average absolute difference (“error”) 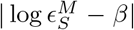 and 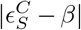 between the true interaction measure *β* and the estimated multiplicative and chimeric epistasis measures, respectively, for all subsets *S* of loci of size |*S*| = *K*. We found (Figure 2B) that the multiplicative measure has a smaller error for all interaction orders *K* and noise parameters *σ*. In particular, we note that the multiplicative measure has smaller error than the chimeric measure even for pairwise interactions with no noise (*K* = 2, *σ* = 0) – i.e., when both the multiplicative and chimeric measures have the same sign – and that the error of the chimeric measure *ϵ*^*C*^ increases with the interaction order *K*. We hypothesize that the reason why the pairwise chimeric measure has much larger error than the pairwise multiplicative measure is because the chimeric measure 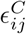 is approximately equal to the (log-)multiplicative measure *only when f*_*ij*_ ≈ 1 and *f*_*i*_*f*_*j*_ ≈ 1, with the two measures being noticeably different otherwise (Figure S2, Supplementary Text). We also emphasize that when there is no noise, i.e. *σ* = 0, the multiplicative measure has zero error, i.e. log *ϵ*^*M*^ = *β*, matching our theoretical results (Theorem 1, Methods).

Our results demonstrate that the multiplicative measure *ϵ*^*M*^ yields a more accurate measurement of pairwise and higher-order epistasis in a multiplicative fitness model compared to the chimeric measure *ϵ*^*C*^ which conflates additive and multiplicative factors.

#### 2.4.2 NK fitness model

We next compared the multiplicative and chimeric epistasis measures using the NK model, a classical model for simulating random fitness landscapes **f** with varying degrees of “ruggedness” [85]. The NK model has two parameters: the number *L* of loci^1^; and *K*, a measure of the ruggedness of the fitness landscape **f**, where the fitness landscape is smoothest at *K* = 0 and most rugged for *K* = *L*− 1. Each locus *𝓁* = 1, …, *L* interacts with *K* random other loci, meaning that the fitness landscape contains at most (*K* + 1)-way interactions. We use the NK model implementation from [49], which simulates fitness values under an additive model, and then exponentiated the NK fitness values to obtain fitness values following a multiplicative model.

Each simulated fitness landscape **f** has an associated graph *G* = (*V, E*) which describes a (simulated) *genetic interaction network*, where the vertices *V* = {1, …, *L*} are the *L* loci and the edges *E* connect pairs of interacting loci [86, 87]. For example, for *K* = 0, the graph *G* has no edges, indicating that there are no interactions between loci, while for *K* = 1, the edges of the graph *G* connect loci with pairwise interactions. (For *K* ≥ 2, one may also describe the interaction relationships with a *hypergraph* where hyperedges connect sets of interacting loci, e.g. [86, 88].)

We find that the chimeric measure falsely indicates the presence of higher-order interactions that are not present in the simulated fitness landscape **f** while the multiplicative measure does not. For example, when the fitness landscape **f** contains only *pairwise* interactions (i.e. *K* = 1), then the 3-way multiplicative epistasis measure 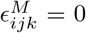 is equal to zero for all triples (*i, j, k*) of loci. However, if the NK model graph *G* contains a triangle (*i, j, k*), then the 3-way chimeric measure 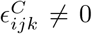 will be nonzero with high probability (Figure 3A). Thus, the chimeric measure *ϵ*^*C*^ falsely indicates the presence of three-way interactions that do not exist in the simulated fitness landscape^2^. More generally, for any value *K >* 0 of the ruggedness parameter, the fitness landscape **f** only contains at most (*K* + 1)- way interactions. The (*K* + 2)-way multiplicative measure *ϵ*^*M*^ is always equal to zero, reflecting that there are no (*K* + 2)-way interactions. However, we empirically observe that the (*K* + 2)-way chimeric measure *ϵ*^*C*^ is often non-zero (Figure 3B).

**Figure 3:**
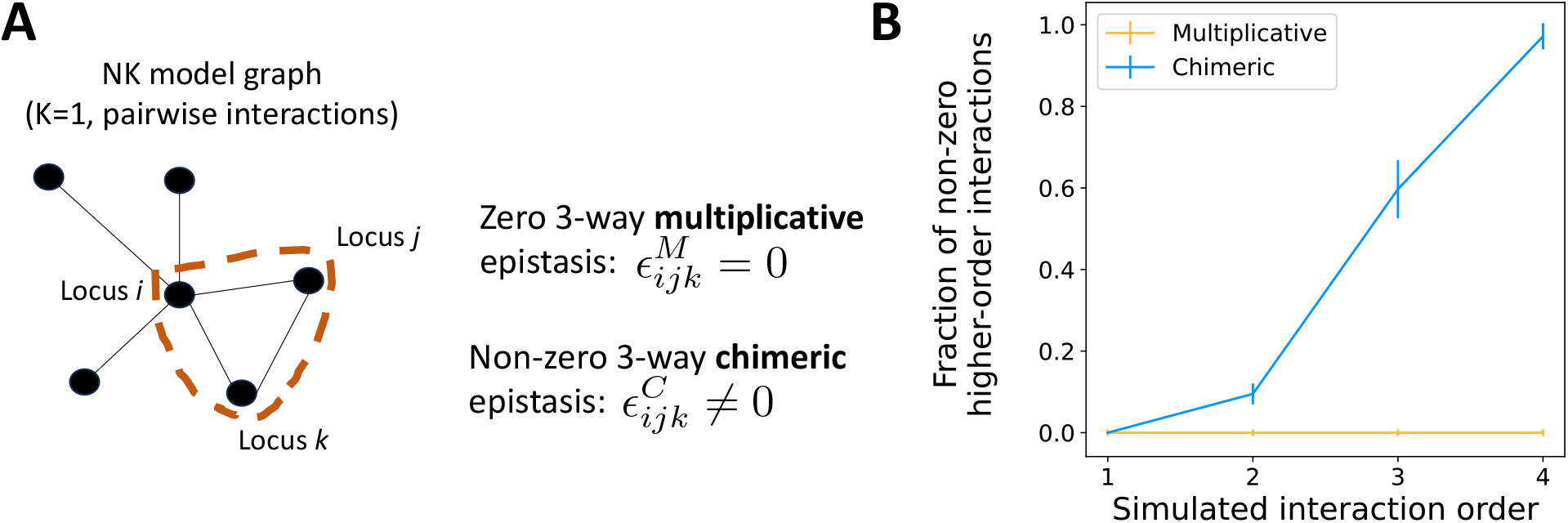
**(A)** A fitness landscape **f** simulated following the NK fitness model with “ruggedness” parameter *K* = 1 contains only pairwise interactions. These interactions are represented with an interaction graph *G*. The 3-way multiplicative measure 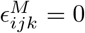 equals zero for all loci triples (*i, j, k*). However, if the triple (*i, j, k*) forms a triangle in the graph *G*, then the 3-way chimeric epistasis measure 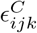 is non-zero with high probability, and incorrectly indicates the presence of a higher-order interaction. **(B)** The fraction of non-zero (*K* + 2)-way interactions (“higher-order interactions”) identified by the multiplicative measure *ϵ*^*M*^ (orange) and the chimeric measure *ϵ*^*C*^ (blue) across 100 fitness landscapes **f** simulated according to the NK fitness model with ruggedness parameter *K*. The fitness landscape **f** contains at most (*K* + 1)-way interactions, but the chimeric measure *ϵ*^*C*^ spuriously detects many non-zero (*K* + 2)-way interactions.

Thus, our analyses demonstrate how the chimeric measure *ϵ*^*C*^ will often erroneously identify higher-order interactions that are not present in the underlying fitness landscape.

### 2.5 Three-way epistasis in budding yeast

We investigate the biological implications of using the chimeric epistasis measure instead of the multiplicative epistasis measure by reanalyzing two triple-gene-deletion studies in budding yeast by Kuzmin et al. [2, 42]. These studies used triple-mutant synthetic genetic arrays (SGA) [90, 91] to measure the fitness of single-, double-, and triple-mutant strains. The authors use a multiplicative fitness model since the SGA protocol models yeast colony sizes as a product of fitness, time, and experimental factors [33]. The Kuzmin et al. studies, [2] and [42], measure fitness values for 195,666 and 256,861 gene triplets, respectively. They calculate the three-way chimeric epistasis measure 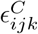 and report 3,196 [2] and 2,466 [42] negative three-way epistatic interactions, respectively.

We calculated the multiplicative epistasis measure 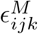 (formula (4)) and the chimeric epistasis measure *ϵ*^*C*^ (formula (5)) used by Kuzmin et al. [2, 42] for the 189,340 gene triplets (*i, j, k*) whose single-, double- and triple-mutant fitness values were available in the publicly available data from [2, 42] and with a reported *p*-value of *p*_*ijk*_ *<* 0.05. Following [2, 42] we say a gene triplet (*i, j, k*) has a *positive chimeric interaction* if 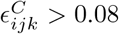; a *negative* chimeric interaction if 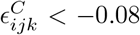 ; and an *ambiguous* chimeric interaction if 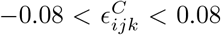. Accordingly, using the same quantile as the chimeric threshold of 0.08, we say that a gene triplet (*i, j, k*) has a *positive (resp. ambiguous, negative) multiplicative interaction* if 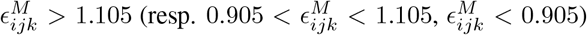. See Supplementary Text for specific details on data processing.

We observed considerable differences between the signs of the multiplicative epistatic measure versus the chimeric epistatic measure (Table 2). In particular, approximately 12% of gene triplets have a *different* interaction sign with the multiplicative measure compared to the chimeric measure. The difference between the two measures is especially pronounced for *negative* interactions, which are typically more functionally informative than positive interactions [2, 42, 33]. In particular, there were 476 gene triplets (*i, j, k*) with a negative *multiplicative-only* interaction, or triplets with a negative multiplicative interaction but not a negative chimeric interaction (Figure 4A). On the other hand, there were only 91 gene triplets with a negative *chimeric-only* interaction, or triplets with a negative chimeric interaction but not a negative multiplicative interaction (Figure 4A); in fact, some of these 91 triplets even had *positive* multiplicative interaction (Figure 4A). We also observe a qualitatively similar discrepancy between the two formula using the earlier fitness data from Kuzmin et al. 2018 [2]; on this data, we find that there were 746 gene triplets with a negative multiplicative-only interaction versus 177 triplets with a negative chimeric-only interaction (Figure S7). Our results were also qualitatively similar when we did not restrict to triplets with reported p-value *p*_*ijk*_ *<* 0.05 (Figure S8).

**Table 2:** Comparison of signs of trigenic interactions in budding yeast calculated using the multiplicative epistasis measure and the chimeric epistasis measure on fitness data from [42]. Red highlighted boxes correspond to gene triplets having different sign of epistasis using the multiplicative measure versus the chimeric measure (approximately 12% of triplets).

| | | Chimeric measure $\epsilon_{ijk}^C$ | | |
| --- | --- | --- | --- | --- |
|  |  | Positive | Ambiguous | Negative |
| Multiplicative measure $\epsilon_{ijk}^M$ | Positive | 1197 | **259** | **0** |
|  | Ambiguous | **116** | 4291 | **91** |
|  | Negative | **10** | **466** | 1527 |

**Figure 4:**
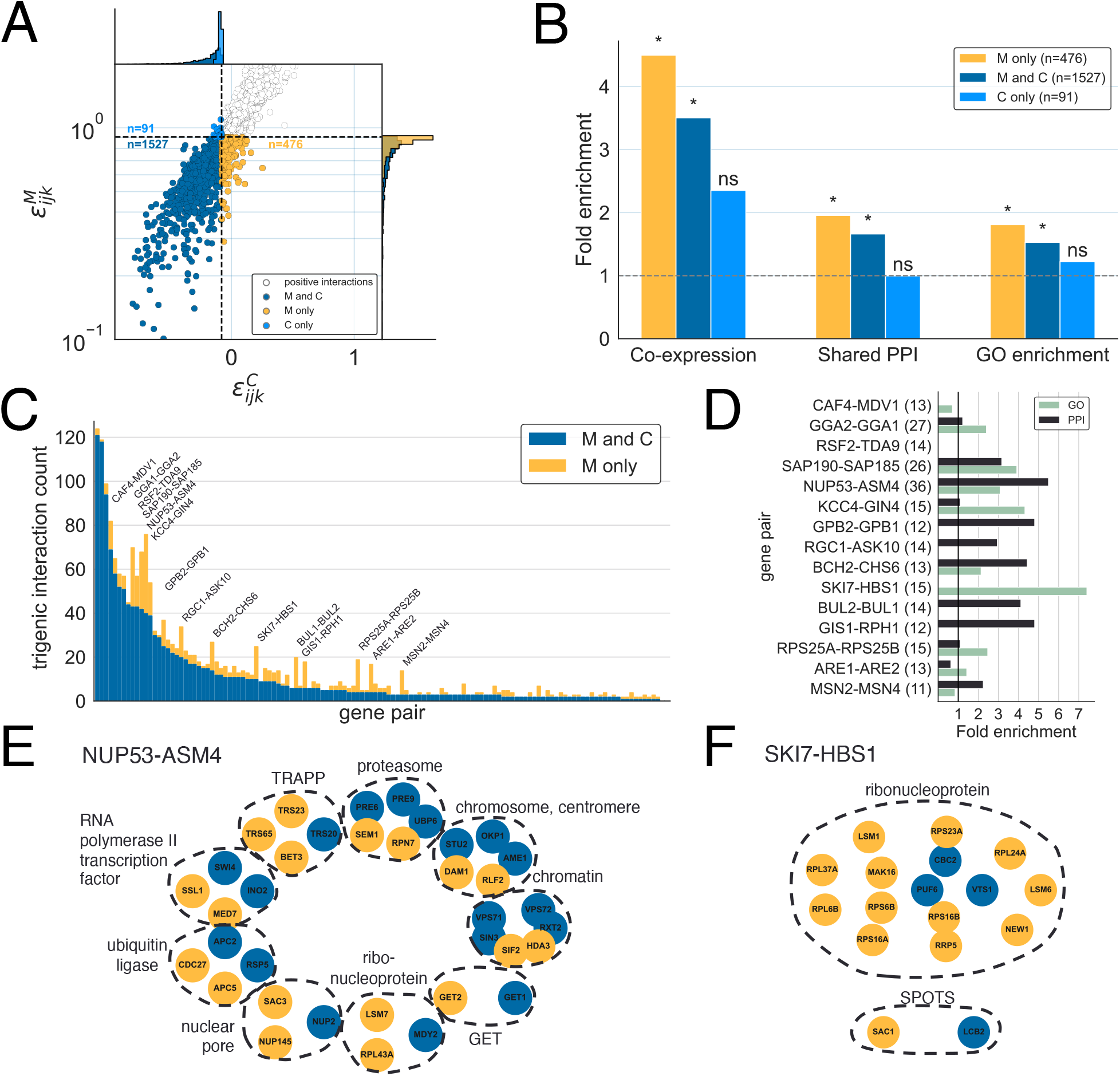
Comparison of negative trigenic interactions in budding yeast calculated using the multiplicative epistasis measure (formula (4)) and the chimeric epistasis measure (formula (5)). **(A)** Chimeric epistasis measure 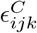 versus the multiplicative epistasis measure 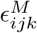 for gene triplets (*i, j, k*) in [42]. We highlight trigenic interactions that are negative only by the multiplicative measure (“M only”), only by the chimeric measure (“C only”), or by both measures (“M and C”). **(B)** Fold enrichment for co-expression patterns, shared protein-protein interactions (PPI), and shared GO annotations for negative trigenic interactions. Asterisk (*) denotes statistical significance at the *P <* 0.05 level, while ‘ns’ indicates not significant. **(C)** Number of negative trigenic interactions (*i, j, k*) for every pair (*i, j*) of genes with at least five negative trigenic interactions. **(D)** Fold enrichment for GO annotations and protein-protein interactions (PPI) for negative “M only” trigenic interactions that involve the gene pairs highlighted in (C). The numbers in parentheses are the number of “M only” interactions. **(E/F)** Genes that have a negative trigenic interaction with either NUP53-ASM4 (E) or with SKI7-HBS1 (F), organized into protein complexes and colored by whether the trigenic interaction is “M only” (gold) or “M and C” (blue).

Negative trigenic interactions often contain genes whose proteins are partially redundant in their functions [92] and are enriched for other features that arise from biological models of functional redundancy, including shared expression patterns [93, 94], shared protein-protein interactions [56], GO annotation, and amino acid divergence [56, 94]. We observed (Figure 4B) that gene triplets with negative multiplicative-only interactions — that is, gene triplets not identified by the chimeric formula used in Kuzmin et al. [42] — are significantly enriched for co-expression (*P* = 0.017, hypergeometric test), shared protein-protein interactions (*P <* 1.5*×*10^−4^, hypergeometric test), and similar GO annotations (*P <* 2.1*×*10^−5^, hypergeometric test). In contrast, gene triplets with a negative chimeric-only interaction are not significantly enriched for any of these features (Figure 4B). In this way using the multiplicative measure extends the network of functionally redundant genes by almost 25% compared to the chimeric measure. We obtain a similar result when analyzing the fitness data from the earlier Kuzmin et al. 2018 study [2] (Figure S7) and also when we do not remove gene triplets with large reported *p*-values *p*_*ijk*_ as computed by [2, 42] (Figure S8). These results demonstrate that using the appropriate three-way multiplicative formula for a multiplicative fitness model leads to more biologically meaningful higher-order genetic interactions compared to using the chimeric epistasis formula that mixes additive and multiplicative scales in an statistically unsound manner.

In particular, trigenic interactions also reveal the functional redundancy of *paralogs*, or pairs of duplicated genes with overlapping functions, since two functionally similar genes tend to have a negative trigenic interaction with a third gene more often compared to gene pairs with non-overlapping functions [42]. Thus, we evaluated whether the gene triplets with negative multiplicative-only interactions involve functionally redundant gene pairs. We quantified the functional redundancy between two genes by calculating the number of negative trigenic interactions to which both genes belong, where we restricted our calculation to gene pairs involved in at least two negative multiplicative interactions. We found that many pairs of genes had additional multiplicative-only interactions (Figure 4C). Thus the multiplicative measure identified additional functional redundancies not found using the chimeric measure. As additional validation, we note that Kuzmin et al. [42] quantify functional redundancy between two genes using a related quantity that they call the *trigenic interaction fraction* (see Supplementary Text for more details). We observed that for most gene pairs, the trigenic interaction fraction is larger when computed using the multiplicative formula versus using the chimeric formula (Figure S10). This observation further supports the conclusion that the multiplicative formula uncovers additional functional redundancies between these paralogs that was not detected by the chimeric measure.

We expect paralogs with large increases in the number of multiplicative-only interactions to be functionally redundant. Of the 130 paralogs we analyzed, there are fifteen paralogs with at least 10 negative multiplicative-only interactions (highlighted in Figure 4C). The three paralogs with the largest number of negative multiplicative-only interactions were RPS24A-RPS25B, MSN2-MSN4, and ARE1-ARE2. For these three paralogs, the multiplicative formula quadrupled the number of total trigenic interactions compared to the number of such interactions reported by [42] using the chimeric formula. These three paralogs also appear to have redundant functions according to other patterns of sequence evolution: all three have highly correlated position-specific evolutionary rates (Table S12 in [42]) and two of them (RPS24A-RPS25B and ARE1-ARE2) have low sequence divergence rates (Figure S9). Moreover, negative genetic interactions have been previously documented for MSN2-MSN4 [95, 96, 97], ARE1-ARE2 [98, 99], and RPS25A-RPS25B [100, 31].

The paralogs with many multiplicative-only interactions are also enriched for shared PPIs or GO annotations with the genes they interact with (Figure 4D). In particular, the paralogs NUP53-ASM4, which are components of the large nuclear pore complex [101], had 36 additional negative multiplicative-only interactions. These epistatic interactions are highly enriched for shared PPIs and GO annotations (Figure 4D) and also involve members of the same protein complexes (Figure 4E). One of the 36 additional genes that interact with NUP53-ASM4 is NUP145, which also forms part of the nuclear pore [102]. Interestingly, while the gene triplet NUP53-ASM4-NUP145 has a negative multiplicative interaction (*ϵ*^*M*^ = 0.684 *<* 1), the same gene triplet was reported to have a *positive* chimeric interaction (*ϵ*^*C*^ = 0.25 *>* 0; [42]). Another example of one of the 36 additional interactions is SAC3, which encodes a nuclear pore-associated protein that functions in mRNA transport [103]. The gene triplet NUP53-ASM4-SAC3 has an extremely negative multiplicative interaction (*ϵ*^*M*^ = 0.046 *<<* 1), but in the original study [42] was reported to have a slightly positive chimeric interaction (*ϵ*^*C*^ = 0.014 *>* 0). Moreover, both NUP145 and SAC3 share at least one protein-protein interaction and GO category with NUP53 and ASM4. These findings provide additional support to the hypothesis by [42] that NUP53 and ASM4 have overlapping functions.

Two other noteworthy paralogs are SKI7 and HBS1; both genes recognize ribosomes stalled during translation and also initiate mRNA degradation. While some studies report that these paralogs have evolved distinct functions [104, 105], other studies show that they retain some overlapping functions [106, 107, 108] and may bind to similar sites on the ribosome [108]. Kuzmin et al. [42] previously reported relatively few (13) trigenic interactions involving both SKI7 and HBS1 as corroboratory evidence for the functional divergence of these paralogs. However, by using the multiplicative epistasis formula, we find 15 additional trigenic interactions involving SKI7 and HBS1. These 15 multiplicative-only interactions are highly enriched for shared GO terms (Figure 4D). Moreover, 12 of the 15 multiplicative-only interactions involve functionally similar genes that are all members of the ribonucleoprotein complex (Figure 4F). Thus, the multiplicative epistasis measure finds evidence for additional functional redundancy between SKI7 and HBS1 that went undetected by the chimeric epistasis measure used in Kuzmin et al. [42].

In addition to the negative interactions just described, we also highlight an example of a biologically relevant *positive* trigenic interaction that is missed by the chimeric epistasis measure but detected by the multiplicative measure. The gene triplet CIK1-VIK1-SUP35td, which consists of two paralogs, CIK1 and VIK1, involved in mitosis [109], and the essential gene SUP35 [110], has an ambiguous, negative chimeric interaction 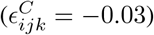 but has an extremely large, positive multiplicative interaction 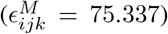. Examining the fitness values (Figure S6) shows that the fitness of the CIK1-VIK1-SUP35td triple mutant is more than 100 times larger than the fitness of the CIK1-SUP35td double mutant. Moreover, positive interactions have been previously documented between pairs of these genes: VIK1 deletion mutants suppress several phenotypes of CIK1 deletion mutants, including a mitotic delay phenotype and a temperature-dependent fitness defect [109]; and a phenotypic suppression interaction exists between CIK1 and SUP35, where deletion of CIK1 reduces the ability of SUP35 to form prions [109]. These previously identified positive pairwise interactions, together with the large triple-mutant fitness value, demonstrate that the gene triplet CIK1-VIK1-SUP35td is more likely to have a positive interaction as indicated by the multiplicative measure, rather than a neutral interaction as indicated by the chimeric measure.

Overall, our results demonstrate not only the degree to which the multiplicative and chimeric formula may lead to distinct interpretations of fitness data, but also that genetic interactions measured using the multiplicative formula appear to be more consistent with other biological features compared to interactions measured using the chimeric formula.

### 2.6 Higher-order interactions in drug responses

We next reanalyzed a drug response dataset [57] in which three-way, four-way, and five-way interactions between drug combinations were quantified using the chimeric formula. For these data, the authors exposed *Escherichia coli* cultures to between one and five antibiotics (out of eight total) at one of three different concentrations. They measured fitness as the difference in exponential growth rates between the culture exposed to antibiotics and a negative control with no antibiotics. The authors then used the chimeric epistasis measure *ϵ*^*C*^ to identify third-, fourth-, and fifth-order interactions between different combinations of antibiotics. We compared their results with the additive epistasis measure *ϵ*^*A*^. We used the additive measure *ϵ*^*A*^ because, under the standard assumption that antibiotic exposure multiplicatively affects the survival probability of individual cells [78], then antibiotic exposure will have an additive effect on the exponential growth rates of the population of cells [111, 112].

The signs of the chimeric interaction measure *ϵ*^*C*^ and the additive interaction measure *ϵ*^*A*^ disagree for three-way, four-way, and five-way interactions, with the discrepancy between the two measures increasing with the interaction order (Figure 5A), which is consistent with our earlier simulations (Figure 1C). The discrepancy is largest for fifth-order interactions, with approximately 14% of fifth-order interactions having a different sign using the additive measure versus the chimeric measure (Figure 5A).

**Figure 5:**
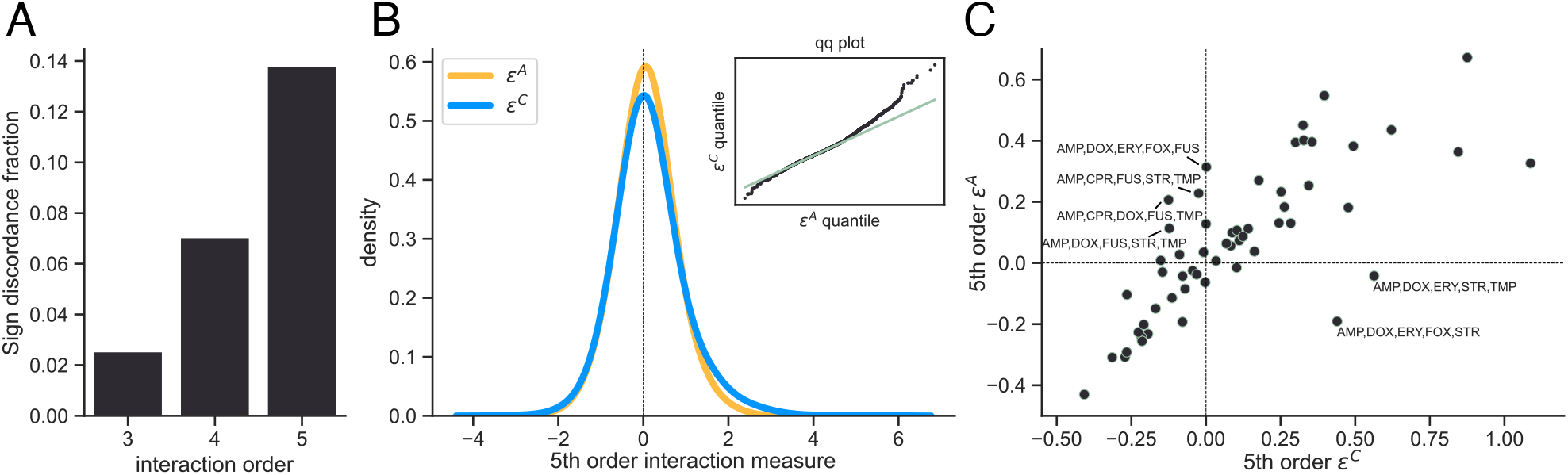
Comparison of the additive and chimeric measures for higher-order interactions between antibiotics in *E. coli* using drug response data from [57]. **(A)** Proportion of *E. coli* cultures where the sign (positive vs. negative) of the chimeric and additive measures disagree. The sign discordance fraction, or proportion of interactions where the sign of the two measures disagree, increases with the interaction order, consistent with the simulations shown in Figure 1. **(B)** Distributions and Q-Q plots (insets) for the additive (orange) and chimeric (blue) measures for 5th order interactions. **(C)** Scatter plot of median relative growth rates for each 5-way combination of antibiotics across concentration levels and replicates.

The discrepancy between the additive and chimeric measures may lead to different conclusions on the type of interactions between antibiotics, i.e. whether a given combination of antibiotics is *synergistic* (more effective at killing bacteria when taken together versus taken individually) or *antagonistic* (less effective together versus individually). For fifth-order interactions, the chimeric measure *ϵ*^*C*^ was more positively skewed than the additive measure *ϵ*^*A*^ (Figure 5B), with a Pearson skewness coefficient of 0.87 for the chimeric measure versus 0.17 for the additive measure. Thus, the chimeric measure is significantly more likely to identify antagonistic interactions than the additive measure (*P <* 7*×*10^−43^, paired t-test).

We then examined specific five-way combinations of antibiotics with different interaction signs following the procedure of [43] and [76]. For each five-way combination of antibiotics we first calculated the median relative growth rate of *E. coli* across replicates and concentrations, and then used these median relative growth values to compute both the additive and chimeric measures (Figure 5C). The interaction between the antibiotic combination Ampicillin (AMP), Doxycycline hyclate (DOX), Erythromycin (ERY), Streptomycin (STR), Trimethoprim (TMP) is highly antagonistic using the chimeric measure (i.e. *ϵ*^*C*^ = 0.56 *>* 0) but synergistic using the additive measure (i.e. *ϵ*^*A*^ = − 0.04 *<* 0). A similar pattern also holds for the antibiotic combination consisting of AMP, DOX, ERY, STR, and Cefoxitin sodium salt (FOX). We emphasize that because we use the same fitness values as reported in [57], the differences between the additive and chimeric measures arise *solely* from the use of the additive versus chimeric measures as opposed to variability arising from biological or technical replicates.

### 2.7 Epistasis between protein mutations

We further demonstrate the difference between the multiplicative and chimeric epistasis measures using experimental fitness data of several different proteins. These fitness values were measured using deep mutational scanning (DMS), a recent class of technologies which use high-throughput sequencing to measure the fitness of many variants of a protein, with fitness measured in terms of the relative frequency of the variant [46]. See Methods for more details on the different datasets. Importantly, recent DMS technologies measure the fitness of molecules with multiple mutations, allowing for the measurement of epistasis between individual mutations. The accurate measurement of epistasis between mutations in proteins is important for many biological applications including 3-D protein structure prediction [115, 68], protein engineering [116], genome editing optimization [114], variant effect prediction [117], and more.

We observe that the multiplicative and chimeric measures have substantial disagreement when measuring higherorder epistasis for several of the proteins. Specifically, we find that the correlation and sign disagreement fraction of the two measures vary as a function of a single quantity: the standard deviation *s* of the 2^*K*^ fitness values {*f*_0···00_, *f*_0···01_ …, *f*_1···11_ } across all *K*-tuples of mutations. In particular, the correlation between the chimeric measure and the multiplicative measure *decreases* as a function of the fitness standard deviation *s* (Figure 6A), while the sign disagreement function *increases* as a function of the fitness standard deviation *s*.

**Figure 6:**
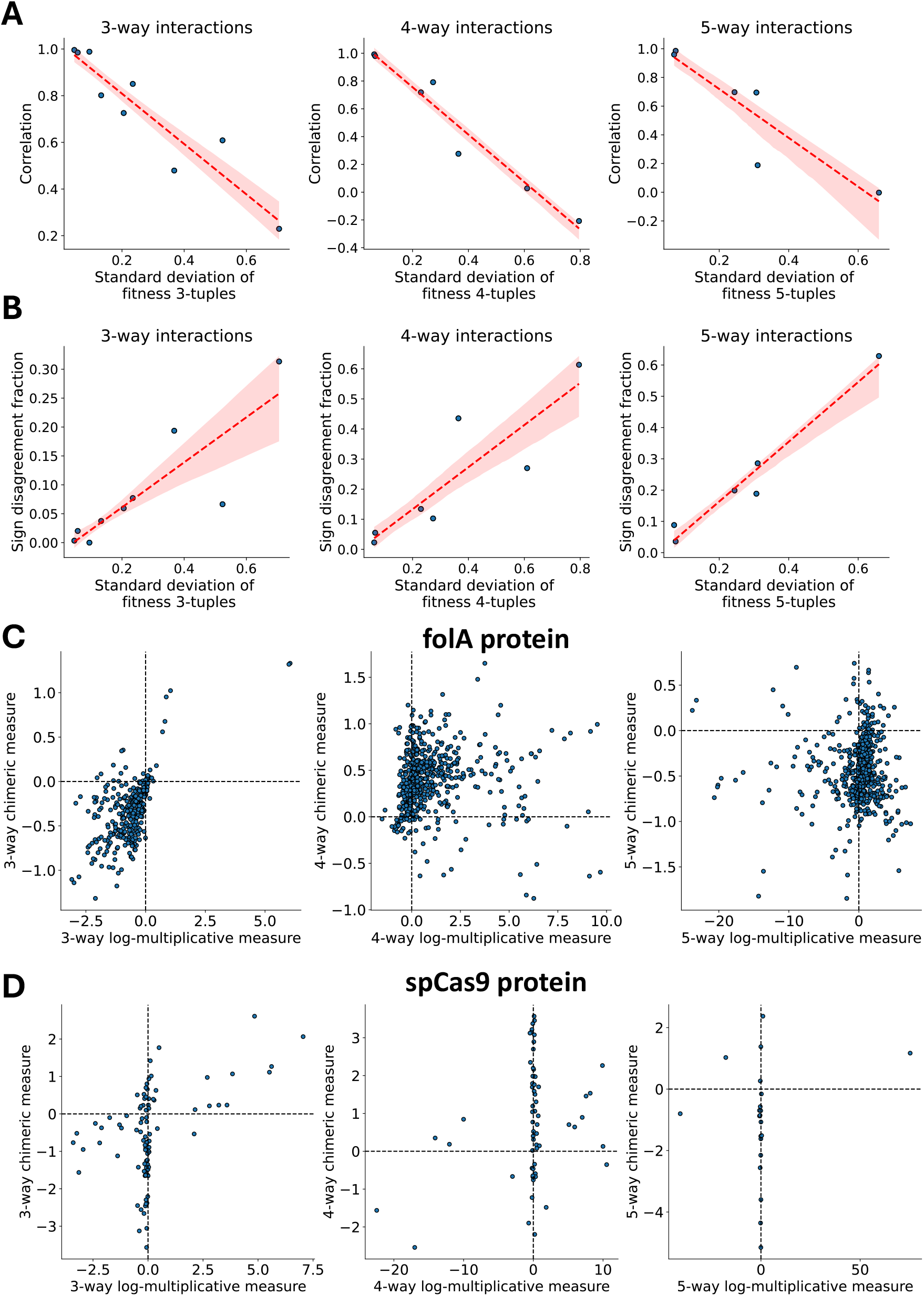
Comparison of multiplicative and chimeric measures for measuring epistasis between protein mutations in nine different proteins. **(A-B)** Standard deviation of fitness values across all (left) three-, (middle) four-, and (right) five-way tuples of mutations versus the average **(A)** correlation and **(B)** sign disagreement fraction of the log-multiplicative measure log *ϵ*^*M*^ versus the chimeric measure *ϵ*^*C*^. **(C-D)** Log-multiplicative measure log *ϵ*^*M*^ versus chimeric measure *ϵ*^*C*^ for the **(C)** *folA* [113] and **(D)** *Streptococcus pyogenes* Cas9 (SpCas9) nuclease [114] proteins.

One protein with a large fitness standard deviation *s* is *folA*, an *E. coli* metabolic protein where the fitness of approximately 260, 000 mutations at nine single-nucleotide loci was recently profiled by [113]. For the folA protein (Figure 6C), the three-way multiplicative and chimeric measures have correlation 0.6086, while the four- and five-way measures have correlation *<* 0.05 — i.e. the two measures are almost uncorrelated for four- and five-way interactions.

There is also substantial sign disagreement between the multiplicative and chimeric measures, with over 60% sign disagreement for five-way interactions. Another biologically meaningful protein with large fitness standard deviation *s* is the *Streptococcus pyogenes* Cas9 (SpCas9) nuclease, a widely used protein for genome editing across biology. The fitness landscape of SpCas9 was profiled by [114], where fitness was measured as the editing efficiency of the SpCas9 protein. For the SpCas9 protein (Figure 6D), the sign disagreement between the two epistasis measures is over 20% for three-, four-, and five-way interactions. The large sign disagreement between the two epistasis measures is likely because for many protein variants, the log-multiplicative measure log *ϵ*^*M*^ is close to 0 while the chimeric measure *ϵ*^*C*^ varies widely between −4 and 2.

Overall, our results demonstrate the extent to which one may infer substantially different higher-order epistasis between protein mutations – including different epistasis *signs* – depending on the measure used.

## 3 Discussion

Higher-order interactions between genetic variants, drugs, and other perturbations play a large role in shaping the fitness landscape of an organism [1, 3, 4, 5]. Yet despite the importance of these interactions, there are multiple different — and sometimes inconsistent — formulae used in the literature for measuring higher-order interactions, most notably for measuring higher-order epistasis between mutations. In particular, many researchers use a *chimeric* formula that quantifies epistasis as an additive deviation from a multiplicative null model and is thus a “chimera” of additive and multiplicative measurement scales.

In this work, we show that there is considerable disagreement between the chimeric epistasis measure and the additive and multiplicative measures. For higher-order interactions, the chimeric measure often has a different *sign* compared to the multiplicative measure (Figure 1C). We demonstrate that this inconsistency is not purely a mathematical curiosity but also leads to markedly different biological conclusions in yeast genetics [2, 42] (Figure 4), antibiotic resistance [57, 76] (Figure 5), and protein epistasis (Figure 6), raising potential questions about some reported higher-order epistatic interactions in the literature. Furthermore, we show that the different epistasis measures are equal to different parametrizations of the multivariate Bernoulli distribution (MVB) [47] (Table 1) and demonstrate that the chimeric epistasis measure is less statistically sound than the additive and multiplicative measures. Our connection between epistasis measures and parameters of the multivariate Bernoulli measure is general and unifies many different epistasis measures: the additive, multiplicative, and chimeric measures; and the Walsh coefficients [50, 41, 49, 60, 69]. Overall, our results demonstrate that the more appropriate multiplicative and additive formula for epistasis yield more mathematically sound and biologically meaningful results compared to the chimeric formula which improperly conflates measurement scales.

To our knowledge, the discrepancy between the chimeric measure and the multiplicative measure has not been previously explored or appreciated in the biological literature. We suspect this is because historically, much of the literature has focused on pairwise interactions, for which the chimeric and multiplicative measures agree on the interaction sign. However, even in the pairwise setting, the two measures have different magnitudes, which may still affect biological findings. For example, Costanzo et al. [31] recently built a large-scale pairwise interaction network for yeast using the chimeric epistasis measure, where they included an edge between two genes if the absolute value of the chimeric measure was greater than a certain threshold. From our results with the trigenic yeast network (Section 2.5), it is possible that the edges in the network would change if one used the more appropriate multiplicative measure instead, which may lead to the inference of different genetic interactions and thus the functional relationships and regulatory mechanisms identified by [31]. As another example, the formulae derived in the theoretical population genetics literature [38, 39, 40] that relate recombination, selection, and the pairwise chimeric epistasis will change if pairwise epistasis is instead measured using the multiplicative measure.

The relevance of higher-order epistasis in human GWAS remains a topic of substantial debate. For example, there are many opinions on whether epistasis is a frequent source of missing heritability for human traits, e.g. [118] argues that epistasis does not contribute to heritability while [119, 120] argue the opposite. We note that several recent papers have identified complex trait epistasis in humans, including pairwise and higher-order epistatic interactions between pathways [121, 122] and individual SNPs [123, 124, 125], suggesting that epistasis is relevant for human genetics.

There are several future directions for our work. First, it would be useful to further investigate the connections between the MVB and regression approaches for higher-order epistasis. For example, regression-based approaches often do not require that one has measured the fitness of all 2^*L*^ genotypes, which may make the estimation of the interaction parameters of the MVB more challenging. Moreover, these regression approaches may sometimes produce biased estimates of epistasis [48], and we imagine that the MVB would provide a useful statistical framework for characterizing such statistical biases. A second direction is to incorporate uncertainty of fitness measurements in the MVB, e.g. by using a Bayesian framework. Thirdly, our statistical framework could be extended to model how higherorder interactions contribute to evolutionary trajectories in a fitness landscape [126, 127, 128]. Fourth, it would be quite interesting to investigate the connections between the MVB and the circuit and Markov bases formulae used to quantify the shape of a fitness landscape [59, 29, 60, 69, 129]. Fifth, for certain technologies, it may be desirable to test whether an additive or multiplicative model better fits experimental fitness data. Our MVB framework provides an approach for doing formal model comparisons. Thus, an interesting and important future direction would be to derive a statistical test using the MVB to test which fitness model better fits data. Finally, one could incorporate environmental, ecological, and other non-genetic factors into the MVB [130, 131].

Ultimately, future studies on interactions in genetics, drug response, protein fitness landscapes, and other domains should take care to use the mathematically appropriate additive or multiplicative formula for measuring higher-order interactions, and not fall victim to the chimera.

## Acknowledgements

We thank Daniel Weinreich, Sriram Sankararaman, and Boyang Fu for helpful comments and suggestions. U.C. was supported by a National Science Foundation Graduate Research Fellowship and the Siebel Scholars program.

B.J.A. gratefully acknowledges financial support from the Schmidt DataX Fund at Princeton University, made possible through a major gift from the Schmidt Futures Foundation. This research is also supported by National Cancer Institute (NCI) grants U24CA248453 and U24CA264027 to B.J.R.

## Authors contributions

Conceptualization: B.J.R., U.C., B.J.A.; methodology: U.C., B.J.R.; software: U.C., B.J.A.; validation: B.J.A.; formal analysis: U.C., B.J.A., and B.J.R.; investigation: U.C., B.J.A., and B.J.R.; data curation: U.C., B.J.A.; writing—original draft: U.C., B.J.A., and B.J.R.; writing—review and editing: U.C., B.J.A., and B.J.R.; visualization: U.C., B.J.A.; supervision: B.J.R.

## Data availability

The datasets used in this study were obtained through publicly available repositories. The data for the analysis in Section 2.5 is available from https://doi.org/10.1126/science.aaz5667 and https://doi.org/10.1126/science.aao1729. The data for the analysis in Section 2.6 is available from https://doi.org/10.1038/s41540-018-0069-9. The data for the analysis in Section 2.7 is described in the Methods section.

## Code availability

The code for the analyses is located in our Github repository.

## 4 Methods

### 4.1 Pairwise epistasis

We start with the simplest setting where the genotype consists of two loci, each with two alleles labeled 0 and 1. Thus, there are four possible genotypes — the wild-type 00, the single mutants 01 and 10, and the double mutant 11 — with corresponding fitness values *f*_00_, *f*_01_, *f*_10_, and *f*_11_ (Figure 1). There are two standard null models that relate genotype to fitness: the additive model and the multiplicative model.

#### Additive fitness model

In the first model, mutations are assumed to have an *additive* effect on fitness [29, 4, 3], e.g. in drug resistance [59, 60, 61, 62, 63] and protein binding [50, 41]. The effect of a mutation is quantified by the difference in fitness when one locus is mutated; for example, *f*_11_ −*f*_10_ measures the effect of a mutation in the second locus, where the genetic background is a mutation in the first locus. An interaction between mutations in the two loci [59, 21], i.e. *pairwise epistasis*, is measured by the difference in the effect of a mutation in one locus across the two possible genetic backgrounds (Figure S1A). The pairwise interaction measure *ϵ*^*A*^ is given by

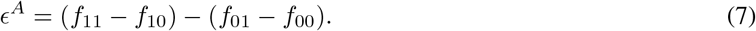

Note that the definition (7) of the pairwise epistasis measure is invariant to the choice of which locus is mutated, i.e. *ϵ*^*A*^ = (*f*_11_ −*f*_10_) − (*f*_01_ − *f*_00_) = (*f*_11_ − *f*_01_) −(*f*_10_ −*f*_00_). In practice, the fitness values are often normalized so that *f*_00_ = 0, i.e. the fitness *f*_00_ of the wild-type is zero, resulting in the following commonly-used equation for pairwise epistasis under an additive fitness model:

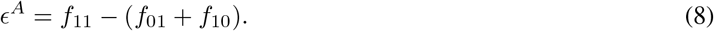

Equivalently, the pairwise epistasis measure *ϵ*^*A*^ is the difference between the observed double-mutant fitness *f*_11_ and the expected double-mutant fitness *f*_01_ + *f*_10_ under a null model with no epistasis. As [4] notes, this definition of pairwise epistasis is similar to Fisher’s original definition of epistasis [132].

The *sign* sgn(*ϵ*^*A*^) of the pairwise epistasis measure *ϵ*^*A*^ determines the type of epistatic interaction. If *ϵ*^*A*^ = 0, then there is no interaction between the two loci and so the fitness *f*_11_ of a double mutant is completely determined by the sum *f*_11_ = *f*_01_ + *f*_10_ of the single mutant fitnesses *f*_01_, *f*_10_. If *ϵ*^*A*^ *>* 0 then there is a *positive* interaction between the two loci, in the sense that the fitness *f*_11_ of the double mutant is *larger* than the fitness if there was no pairwise interaction. Similarly, if *ϵ*^*A*^ *<* 0 then there is a *negative* interaction between the two loci, in the sense that the fitness *f*_11_ of the double mutant is *smaller* than the fitness if there was no pairwise interaction.

The pairwise epistasis measure *ϵ*^*A*^ is equivalent to two other notions of epistasis used in the genetics literature. First, the pairwise epistasis measure *ϵ*^*A*^ is equal to the pairwise interaction term in the standard linear *regression* framework for quantifying epistasis [3, 21]. Specifically, if the fitness values *f*_00_, *f*_01_, *f*_10_, *f*_11_ follow a linear model of the form

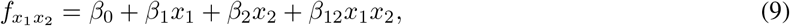

then the coefficient *β*_12_ of the interaction term *x*_1_*x*_2_ is equal to the pairwise epistasis measure *ϵ*^*A*^ in (7). Second, the epistasis measure *ϵ*^*A*^ is equal (up to a constant factor) to the 2nd-order Walsh coefficient that is often used to measure “background-averaged” epistasis [41, 49, 50, 60, 69].

#### Multiplicative fitness model

In this model, mutations are assumed to have a *multiplicative* effect on fitness, e.g. modeling cellular growth rates [4, 23, 33, 2, 31, 65, 66]. The multiplicative pairwise epistasis measure (Figure S1B) is given by

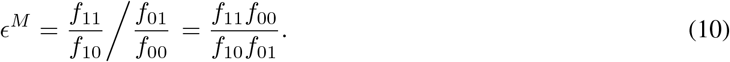

As in the additive model, in practice the fitness values are typically normalized such that *f*_00_ = 1, resulting in the following equation for pairwise epistasis:

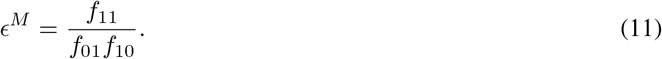

That is, the pairwise epistasis measure *ϵ*^*M*^ is the *ratio* between the double-mutant fitness *f*_11_ and the product *f*_01_*f*_10_ of the single-mutant fitness values.

The multiplicative fitness model is closely related to the additive fitness model: if fitnesses *f* are multiplicative, then the *log-*fitnesses log *f* are additive. Thus, the sign of the interaction is determined by the difference between the epistasis measure *ϵ*^*M*^ and 1, or equivalently the sign sgn(log *ϵ*^*M*^) of the log *ϵ*^*M*^ of the epistasis measure *ϵ*^*M*^ (Figure 1A). If *ϵ*^*M*^ *>* 1, i.e. log *ϵ*^*M*^ *>* 0, there is a *positive* interaction between the two loci; if *ϵ*^*M*^ = 1, i.e. log *ϵ*^*M*^ = 0, then there is no interaction between the two loci; and if *ϵ*^*M*^ *<* 1, i.e. log *ϵ*^*M*^ *<* 0, then there is a *negative* interaction between the two loci.

The multiplicative pairwise epistasis measure is closely related to the pairwise interaction term in the standard *loglinear* regression framework for epistasis [3, 21]. Specifically, if the fitness values *f*_00_, *f*_01_, *f*_10_, *f*_11_ follow a log-linear regression model of the form

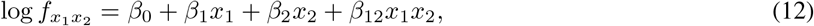

then *β*_12_ = *ϵ*^*M*^.

#### Chimeric formula

Many studies in genetics use a multiplicative fitness model but do not measure pairwise epistasis with the multiplicative epistasis measure *ϵ*^*M*^. Instead, these papers use a multiplicative null model but measure deviations with an *additive* scale, yielding the following epistasis measurement:

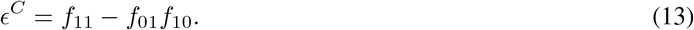

We call 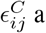 *“chimeric”* measure as it measures deviations from a multiplicative null model on an additive scale, and is thus a *chimera* of both the multiplicative and additive measurement scales. The chimeric measure has been widely used in the genetics literature (e.g. [23, 24, 25, 26, 27, 28, 29, 4, 30, 31, 32, 33, 34]) and in the drug interaction literature (e.g. [70, 71, 72, 73, 74, 75, 57, 43, 76, 77]). In these applications, similar to the additive measure, the sign of an interaction between two loci is determined by the sgn(*ϵ*^*C*^) of the chimeric measure *ϵ*^*C*^: *ϵ*^*C*^ *>* 0 corresponds to a positive interaction while *ϵ*^*C*^ *<* 0 corresponds to a negative interaction.

Although it is often described in terms of a multiplicative fitness model, the chimeric epistasis measure *ϵ*^*C*^ is *not* equal to the multiplicative measure *ϵ*^*M*^. The chimeric epistasis measure *ϵ*^*C*^ in equation (13) is similar to equation (11), but the deviation between the observed double-mutant fitness *f*_11_ and the expected fitness *f*_01_*f*_10_ under a multiplicative null model is computed using subtraction instead of division. Equivalently, the (log-)multiplicative epistasis measure log *ϵ*^*M*^ = log *f*_11_ − log *f*_01_*f*_10_ computes the difference between the observed and expected logarithm of the fitness of the double mutant, while the chimeric epistasis measure *ϵ*^*C*^ = *f*_11_− *f*_01_*f*_10_ computes the difference directly (Figure 1A). In this way, the chimeric epistasis measure may overstate or understate the strength of a pairwise interaction in a multiplicative fitness model (Figure 1A); see Supplementary Text for a numerical example highlighting this issue with the chimeric measure.

Nevertheless, we show that the chimeric measure *ϵ*^*C*^ measures the same *sign* of an interaction as the multiplicative measure *ϵ*^*M*^.

##### Proposition 1.

*Let f*_01_, *f*_10_, *f*_11_ ∈ ℝ *be real numbers. Let* 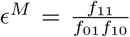 *and ϵ*^*C*^ = *f*_11_ − *f*_01_*f*_10_. *Then* sgn(*ϵ*^*C*^) = sgn(log *ϵ*^*M*^).

*Proof*. 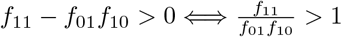.

### 4.2 Higher-order epistasis

We next generalize our discussion to genotypes with *L*≥ 2 loci, where we demonstrate that the differences between the multiplicative measure and the chimeric measure become even more pronounced when analyzing *higher-order epistasis*, or interactions between three or more loci.

There are 2^*L*^ genotypes *x*_1_ · · · *x*_*L*_, where *x*_*𝓁*_ ∈ {0, 1} indicates a mutation in locus *𝓁*, with each genotype *x*_1_ · · · *x*_*L*_ having a corresponding fitness value 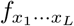, e.g. *f*_010_ is the fitness of genotype 010 with a mutation in the second locus and no mutations in the first and third loci. However, because writing out the 2^*L*^ genotypes is infeasible for large *L*, we use the following notational shorthand. We use *f*_*i*_ to refer to the fitness of the genotype with a single mutation in locus *i, f*_*ij*_ to refer to the fitness of the genotype with mutations in loci *i, j*, and so on. For example, for *L* = 3 loci, *f*_2_ corresponds to *f*_010_ while *f*_12_ corresponds to *f*_110_. Without loss of generality we assume the wild-type fitness *f*_∅_ is equal to 0 for the additive fitness model, and equal to 1 for the multiplicative and chimeric fitness models. We also define 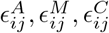 as the additive, multiplicative, and chimeric pairwise epistasis measure, respectively, between the *i*-th locus and the *j*-th locus, i.e. 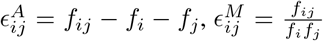 and 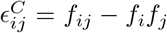. For example, for *L* = 3 loci, 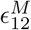 corresponds to 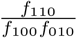.

#### Additive fitness model

We start by quantifying three-way epistasis in the additive fitness model. When there is no pairwise epistasis, the fitness *f*_*ijk*_ of a triple mutant is equal to *f*_*i*_ + *f*_*j*_ + *f*_*k*_, i.e. the fitness from of each of the single-mutants. When there is pairwise epistasis, then the triple mutant fitness *f*_*ijk*_ also includes pairwise interactions measures, i.e.

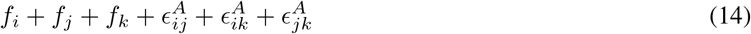

Three-way epistasis is computed by measuring the difference between the observed triple-mutant fitness *f*_*ijk*_ and the expected fitness in (14) when only pairwise interactions are included. Thus, the three-way additive epistasis measure 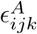 is given by

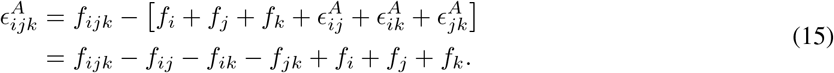

As in the pairwise case, the sign of the three-way epistatic measure 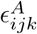 determines the sign of the interaction: if 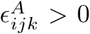, then there is a *positive* three-way interaction between loci *i, j, k* — in the sense that the fitness *f*_*ijk*_ of the triple mutant is larger than the expected fitness in (14) when only pairwise interactions are present — while if 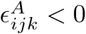, then there is a *negative* three-way interaction between loci *i, j, k*.

Our derivation of the three-way epistasis measure 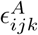 is easily extended to higher-order interactions. The additive *K*-way epistasis measure 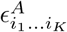 is defined recursively as

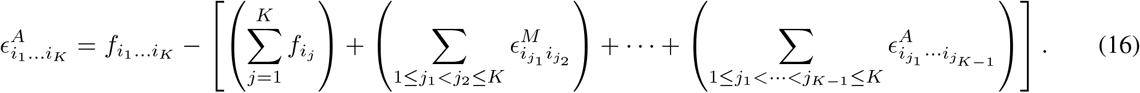

The *K*-way epistasis measures 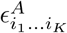 are proportional to two other measures of epistasis: (1) the *K*-th order Walsh coefficient used to quantify background-averaged epistasis among *K* genetic loci [41, 49, 50] and (2) the *K*-th order interaction coefficients of a linear regression model, which we discuss in more detail in Section 4.3.

#### Multiplicative fitness model

We derive formulae for epistasis in a multiplicative fitness model by using the equivalence between multiplicative fitness and additive log-fitness. For example, the 3-way epistasis measure 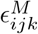 in the multiplicative model is given by

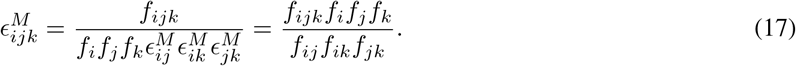

As in the pairwise setting, the sign of interaction is determined by the difference between the multiplicative measure 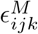 and 1, or equivalently by the sgn 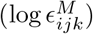of the logarithm of the epistasis measure 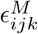.

Using (16), then the *K*-way epistasis measure 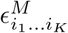 in the multiplicative model is defined recursively by

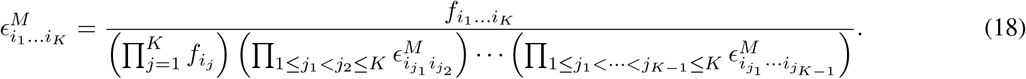

Recent work in the genetics [2, 42] and drug interaction [43] claim to measure three-way epistasis using a multiplicative fitness model. However, they do not measure three-way epistasis the multiplicative epistasis formula (17) but instead derive a chimeric formula using both additive and multiplicative measurement scales:

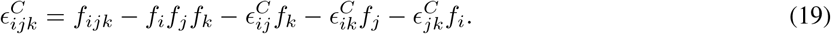

We call 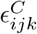 the *chimeric* three-way epistasis measure. In these applications, the sign of the interaction is determined by the sgn 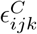 of the chimeric measure 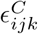.

Despite the claim that the chimeric measure 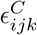 is derived from a multiplicative fitness model, it is clear by inspection that the three-way chimeric measure 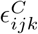 is not equal to the multiplicative three-way epistasis measure 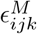.

However, unlike in the pairwise setting, even the *signs* of these two measures disagree (Figure 1B). We demonstrate in the Supplementary Text that even when 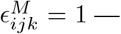 that is, there is no three-way epistasis — the chimeric three-way epistasis measure 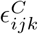 may still indicate either positive or negative three-way epistasis.

Tekin et al. [43] extended the three-way chimeric epistasis formula (19) by heuristically deriving chimeric formulae for 4-way and 5-way epistasis. For example, their chimeric formula for 4-way epistasis is given by

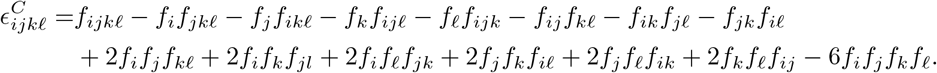

As in three-way epistasis, the sign of the 4-way and 5-way chimeric epistasis measures derived by [43] do not match the signs of the corresponding multiplicative epistasis measures (Figure 1C). This fundamental disagreement motivates a deeper mathematical understanding of these epistasis measures, which we explore in the following section.

### 4.3 Multivariate Bernoulli distribution

In the previous section, we defined quantitative measures of epistasis for two standard null models for fitness: the additive model and multiplicative model. Nevertheless, some recent papers use a multiplicative fitness model but instead use an epistasis measure which is a *chimera* of both multiplicative and additive measurement scales. Here, we unify these different epistasis measures using the *multivariate Bernoulli* distribution from probability theory [47].

The multivariate Bernoulli distribution describes any distribution on {0, 1}^*L*^, i.e. binary strings of length *L*, for *L*≥ 2. The multivariate Bernoulli distribution has three different parameterizations which are used throughout the literature [47, 82]. We start by describing these parametrizations for the simplest such distribution: a *bivariate* Bernoulli distribution over binary strings of length *L* = 2.

#### Bivariate Bernoulli distribution

Suppose that *X* = (*X*_1_, *X*_2_) ∈ {0, 1} ^2^ is distributed according to a bivariate Bernoulli distribution. A distribution on *X* is specified by the parameters *p*_00_, *p*_01_, *p*_10_, *p*_11_, where 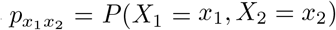 is the probability of (*x*_1_, *x*_2_). The parameters **p** = (*p*_00_, *p*_01_, *p*_10_, *p*_11_) are sometimes called the *general* parameters [47]. Note that since *p*_00_ + *p*_01_ + *p*_10_ + *p*_11_ = 1, only three such parameters are needed to define the distribution.

The probability density function (PDF) *P* (*X*_1_, *X*_2_) of *X* = (*X*_1_, *X*_2_) has the form

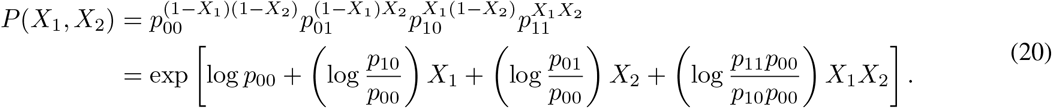

In other words, the PDF *P* (*X*_1_, *X*_2_) follows a log-linear model of the form

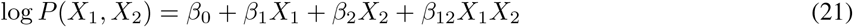

for constants *β*_0_, *β*_1_, *β*_2_, *β*_12_ ∈ ℝ. There is a one-to-one correspondence between the general parameters **p** = (*p*_00_, *p*_01_, *p*_10_, *p*_11_) and the constants ***β*** = (*β*_0_, *β*_1_, *β*_2_, *β*_12_). Thus, a bivariate Bernoulli distribution is also parametrized by the parameters ***β***, also known as the *natural* parameters of the distribution [47]. As with the general parameters **p**, we note that only three out of the four parameters *β*_0_, *β*_1_, *β*_2_, *β*_12_ are needed to fully specify a distribution. We also note that independence between the random variables *X*_1_ and *X*_2_ is described by the parameter *β*_12_, where *X*_1_ and *X*_2_ are independent if and only if *β*_12_ = 0.

Equation (21) demonstrates that *X* = (*X*_1_, *X*_2_) follows an *exponential family* distribution, a wide class of distributions that includes many common distributions including normal distributions or Poisson distributions. In particular, using the terminology of exponential families, equation (21) shows that the *sufficient statistics* of *X* are *X*_1_, *X*_2_, and *X*_1_*X*_2_, with corresponding *canonical* parameters *β*_1_, *β*_2_, and *β*_12_ [83, 133, 134]. As a result, the distribution *P* (*X*) is uniquely defined by the *expected values E*[*X*_1_], *E*[*X*_2_], *E*[*X*_1_*X*_2_] of the sufficient statistics, sometimes called the *moments* or the *mean parameters* of the distribution [83]. Thus, we obtain a third parametrization of the distribution *P* (*X*) using the *moments µ*_0_ = 1, *µ*_1_ = *E*[*X*_1_], *µ*_2_ = *E*[*X*_2_], *µ*_12_ = *E*[*X*_12_]. The elements of the vector ***µ*** = (1, *µ*_1_, *µ*_2_, *µ*_12_) of moments are sometimes called the *mean parameters* of the distribution.

#### Multivariate Bernoulli distribution

The three parametrizations we derived for the bivariate Bernoulli distribution extend to the multivariate Bernoulli distribution. Suppose that (*X*_1_, …, *X*_*L*_) ∈ {0, 1}^*L*^ is distributed according to a multivariate Bernoulli distribution. Then the distribution *P* (*X*) of the random variables *X* is uniquely specified by one of the three following parametrizations.

##### 1. General parameters

These are 2^*L*^ non-negative values 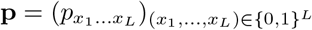 satisfying

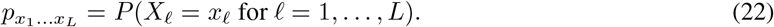

For example if *L* = 3, then *p*_010_ = *P* (*X*_1_ = 0, *X*_2_ = 1, *X*_3_ = 0) and *p*_110_ = *P* (*X*_1_ = 1, *X*_2_ = 1, *X*_3_ =0). Note that since 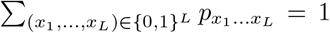, only 2^*L*^ − 1 values 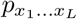 are necessary to define the distribution.

##### 2. Natural/canonical parameters

These are 2^*L*^ real numbers ***β*** = (*β*_*S*_)_*S*⊆[*L*]_ ∈ ℝ satisfying

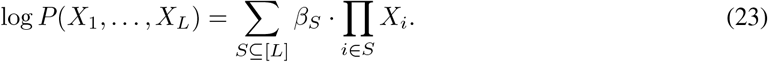

Similar to the general parameters *p*_*i*_, only 2^*L*^ − 1 values *β*_*S*_ are necessary to uniquely define the distribution. Typically, the parameter *β*_∅_, often called a normalizing constant or a *partition function* of the distribution, is left unspecified. As noted in the bivariate setting, equation (23) shows that the multivariate Bernoulli is an exponential family distribution with 2^*L*−1^ sufficient statistics of the form ∏_*i*∈*S*_ *X*_*i*_ for subsets *S* with |*S*| *>* 0.

Moreover, by rewriting (23) as

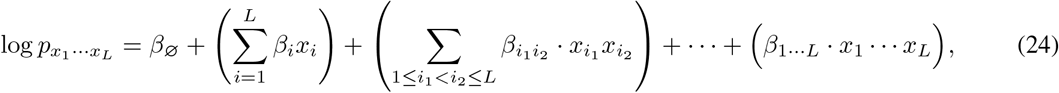

we observe that the natural parameters ***β*** correspond to interaction coefficients in a log-linear regression model with response variables **p**. For example, the natural parameter *β*_12_ is the coefficient of the interaction term *x*_1_*x*_2_.

##### 3. Moments/mean parameters

These are 2^*L*^ real numbers ***µ*** = (*µ*_*S*_)_*S*⊆[*L*]_ satisfying

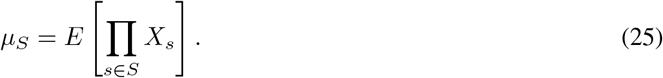

For example if *L* = 3, then *µ*_13_ = *E*[*X*_1_*X*_3_] while *µ*_12_ = *E*[*X*_1_*X*_2_]. The mean parameters {*µ*_*S*_}_|*S*|*>*0_ are sufficient statistics for the multivariate Bernoulli distribution, as seen in the exponential family form (23) of the multivariate Bernoulli distribution.

We note that all three parametrizations, as well as the fitness values *f* and epistasis measures *ϵ*, can be defined either in terms of subsets *S* ⊆ [*L*] as with the natural parameters ***β*** and moments ***µ***, or in terms of binary strings *x*_1_ · · · *x*_*L*_ as with the general parameters **p**. We use both definitions interchangeably, with the convention that a subset *S* ⊆ [*L*] corresponds to the binary string *x*_1_ · · ·*x*_*L*_ with *x*_*i*_ = 1_{*i*∈*S*}_.

Moreover, when written as vectors indexed by binary strings, the three parametrizations ***β, µ*, p** of the multivariate Bernoulli are related to each through different linear transformations involving a matrix operation known as the *Kronecker product* (see Supplementary Text for specific formulae). Interestingly, several papers quantify epistasis using the *Walsh-Hadamard transform* which is also defined in terms of Kronecker products [41, 49, 50]. This connection is not a coincidence; in the next section we show that the Walsh-Hadamard transform is closely related to the parametrizations of the multivariate Bernoulli.

### 4.4 Unifying epistasis measures with the multivariate Bernoulli

The multivariate Bernoulli distribution provides an elegant means by which to describe the different epistasis formulae in the literature. We model the genotype (*X*_1_, …, *X*_*L*_) ∈ {0, 1}^*L*^ as a random variable distributed according to a multivariate Bernoulli distribution. The parametrizations of the multivariate Bernoulli correspond to different features of the genotype; for example, the probability 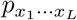 might be derived from the frequency of observing the genotype (*x*_1_, …, *x*_*L*_) in a large population.

#### 4.4.1 Multiplicative and additive epistasis measures

We start by relating the multiplicative epistasis formula (18) to the multivariate Bernoulli distribution. A careful reader may observe that the natural parameter 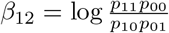 in the bivariate Bernoulli distribution (21) bears close resemblance to the multiplicative epistasis measure in equation (10). Specifically, if the fitness 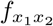 of each genotype (*x*_1_, *x*_2_) ∈ {0, 1}^2^ is proportional to the probability 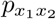 of that genotype in the multivariate Bernoulli, then the natural parameter *β*_12_ is equal to the logarithm log 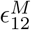 of the multiplicative epistasis measure 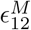. Thus, for *L* = 2 loci, epistasis is measured by the natural parameters ***β*** of a bivariate Bernoulli distribution.

We prove that this observation is not specific to the bivariate Bernoulli distribution with *L* = 2 loci, and in fact generalizes to any number *L* of loci. Specifically, we prove that if the fitness 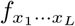 of genotype (*x*_1_, …, *x*_*L*_) is proportional to the probability 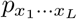 of observing the genotype, then for each subset *S* ⊆ [*L*] of loci, the natural parameter *β*_*S*_ equals the logarithm log 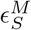 of the corresponding multiplicative epistasis measure as defined in equation (18).

##### Theorem 1.

*Let f*_x_ ∈ℝ *be fitness values for genotypes* **x** = (*x*_1_, …, *x*_*L*_) ∈ {0, 1} ^*L*^ *such that f*_x_ = *c* · *p*_x_ *for some constant c >* 0 *and for some multivariate Bernoulli random variable* (*X*_1_, …, *X*_*L*_) *with general parameters* 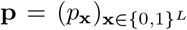. *Then for all subsets S* ⊆ {1, …, *L*} *of loci, the log multiplicative epistasis measure* log 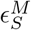 *is equal to the interaction parameter β*_*S*_ *of the random variable* (*X*_1_, …, *X*_*L*_).

By using the equivalence between multiplicative fitness values and additive log-fitness values, we also derive a similar probabilistic interpretation of the additive epistasis formula. Specifically, if fitness 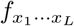 is proportional to the *log*-probability log 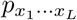 of observing the genotype (*x*_1_, …, *x*_*L*_), then for each subset *S* = {*i*_1_, …, *i*_*K*_} ⊆ [*L*] of loci, the natural parameter *β*_*S*_ equals the logarithm 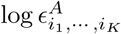 of the corresponding additive epistasis measure as defined in equation (16). We formalize this observation as the following Corollary of Theorem 1.

##### Corollary 1.

*Let f*_x_ ∈ℝ *be fitness values for genotypes* **x** = (*x*_1_, …, *x*_*L*_) ∈ {0, 1} ^*L*^ *such that f*_x_ = *c* · log *p*_x_ *for some constant c >* 0 *and for some multivariate Bernoulli random variable* (*X*_1_, …, *X*_*L*_) *with general parameters* 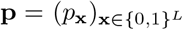. *Then for all subsets S* ⊆ {1, …, *L*} *of loci, the log additive epistasis measure* log 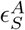 *is equal to the interaction parameter β*_*S*_ *of the random variable* (*X*_1_, …, *X*_*L*_).

The assumption that the fitness 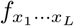 of a genotype (*x*_1_, …, *x*_*L*_) is proportional to the probability 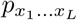 of observing the genotype is often used in generative models for estimating the fitness of protein structures from sequence data [80, 81]. This assumption also has a natural biological interpretation: suppose the fitness value *f*_x_ corresponds to the growth rate of an organism with genotype **x** = (*x*_1_,…, *x*_*L*_), and suppose that initially there are an equal number of organisms of each of the 2^*L*^ genotypes **x** ∈ {0, 1} ^*L*^. Then after one unit of time, the frequency *p*_x_ of each genotype **x** will be proportional to its growth rate *f*_x_.

We also note that the statistical problem of estimating the natural parameters ***β*** or mean parameters ***µ*** of a multivariate Bernoulli distribution from samples (*X*_1_, …, *X*_*L*_) of the distribution is computationally hard [83]. The reason why we are able to use relatively simple formulae (16), (18) to compute the natural parameters ***β*** is because in this setting, we have both samples (*X*_1_, …, *X*_*L*_) *and* their corresponding probabilities *P* (*X*_1_, …, *X*_*L*_), i.e. the fitness values *f*.

##### Relationship with (log-)linear regression

Under the assumption that the fitness values 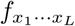 are proportional to the genotype probabilities 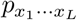, then (24) is a log-linear regression model of the form

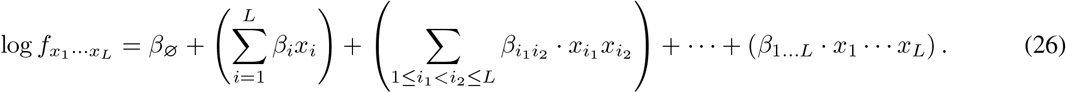

Thus, Theorem 1 shows that computing the multiplicative epistasis measure *ϵ*^*M*^ is equivalent to computing the interaction parameters of the log-linear regression in (26). The interaction parameters of a regression are a standard approach for quantifying epistasis in GWAS and eQTL analyses [3].

Similarly, Corollary 1 demonstrates the equivalence between the additive epistatic measure *ϵ*^*A*^ and the coefficients of a linear regression model with response variables equal to the fitness values. Specifically, under the assumption that the fitness values 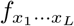 are proportional to the logarithm log 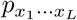 of the genotype probabilities, then computing the additive epistasis measures *ϵ*^*A*^ is equivalent to computing the interaction parameters ***β*** of the following linear regression model

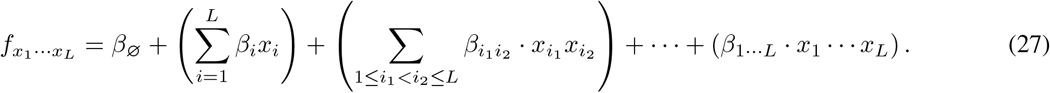

In this way, Theorem 1 and Corollary 1 provide a connection between the multiplicative and additive epistasis measures and the interaction coefficients of log-linear and linear regression models, respectively.

##### Relationship with case-control GWAS

We also prove that the natural parameters ***β*** of a MVB with three variables are closely related to the two standard approaches for measuring pairwise SNP-SNP interactions in a case-control GWAS: logistic regression and conditional independence testing [84]. Specifically, suppose we are given genotype (*X*_1_, *X*_2_) ∈ {0, 1} ^2^ and (binary) disease status *D* ∈ {0, 1}. Then the joint random variable (*X*_1_, *X*_2_, *D*) follows a MVB distribution, where the log-probability log *P* (*X*_1_, *X*_2_, *D*) is given by the following expression in terms of the natural parameters ***β***:

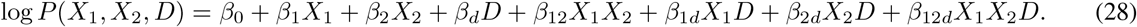

Note that there is a natural approach for representing GWAS data from diploid genomes (with {0, 1, 2} -valued allelic states) using binary random variables *X*_1_, *X*_2_, as described in [135].

We show that the logistic regression approach for measuring pairwise interactions is equivalent to computing *β*_12*d*_, while the conditional independence test is equivalent to testing the null hypothesis *H*_0_ : *β*_12_ = *β*_12*d*_ = 0. See the Supplementary Text for proofs.

#### 4.4.2 Chimeric epistasis measure

The multivariate Bernoulli also provides a way of rigorously defining the pairwise and higher-order chimeric epistasis measures using *joint cumulants*. Joint cumulants are a concept from probability theory used to quantify higher-order interactions between random variables. For example, the 2nd order joint cumulant *κ*(*X, Y*) of two random variables *X, Y* is given by

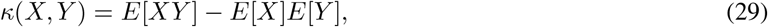

and is equal to the covariance Cov(*X, Y*). The 3rd order joint cumulant *κ*(*X, Y, Z*) of three random variables is given by

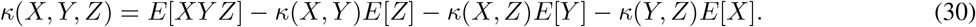

Under the assumption that the fitness 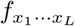 of a genotype (*X*_1_, …, *X*_*L*_) is equal to the corresponding *moment* 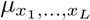, we define the *K*-way chimeric epistatic measure 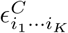 as the *K*-th order *joint cumulant* 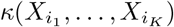 of the random variables 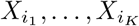.

##### Definition 1.

*Let f*_x_ ∈ ℝ *be fitness values for genotypes* **x** = (*x*_1_, …, *x*_*L*_) ∈ {0, 1}^*L*^ *such that* 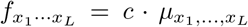 *for some constant c >* 0 *and for some multivariate Bernoulli random variable* (*X*_1_, …, *X*_*L*_) *with moments* 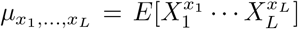.*The* chimeric epistasis measure 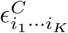 *is the joint cumulant* 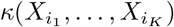 *of the random variables* 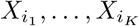.

Our definition of the *K*-th order chimeric epistasis measure 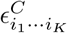 as the *K*-th order joint cumulant formalizes the heuristic derivation of the chimeric measure in previous literature. Almost every paper that uses the chimeric epistasis measures *ϵ*^*C*^ does not even mention the joint cumulant. Two notable exceptions are [57, 76], which use the joint cumulants to derive formulae for 3-way, 4-way, and 5-way interactions between drugs. However, [57, 76] do not rigorously define a probability distribution nor the random variables whose joint cumulant they compute.

At the same time, our formal definition of the chimeric epistasis measure *ϵ*^*C*^ reveals two critical issues with the chimeric formula. First, the assumption that the fitness values *f* are equivalent to the moments of a MVB random variable is not biologically reasonable for higher order interactions between three or more loci. This assumption implies that the fitness of a particular genotype depends on the probability of many other genotypes. For example, making this assumption for *L* = 4 loci, the fitness *f*_1100_ of a double mutant is equal to the moment *E*[*X*_1_*X*_2_], which is equal to

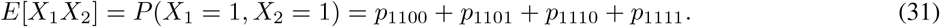

However, it is not clear why the fitness *f*_1100_ of a *single* genotype, 1100, should equal depend on the probabilities of *four different* genotypes, 1100, 1101, 1110, and 1111.

The second issue is that joint cumulants are not necessarily an appropriate measure of higher-order interactions between *binary* random variables. The differences between the joint cumulants and natural parameters ***β*** have been previously investigated in the neuroscience literature, as both quantities have been used to quantify higher-order interactions in neuronal data. For example, Staude et al. [54, 55] write that the joint cumulants *κ* and natural parameters ***β*** measure mathematically distinct types of higher-order interactions, and that each quantity may be appropriate for different applications. In particular, Staude et al. note that the joint cumulants measure higher-order interactions between random variables in terms of “additive common components”, while the natural parameters ***β*** measure *“to what extent the probability of certain binary patterns can be explained by the probabilities of its sub-patterns”*. It follows that for binary mutation data, the natural parameters ***β*** correspond exactly with the epistasis we aim to measure, i.e. how the fitness of a binary pattern can be explained by the fitness of its “sub-patterns”, while the joint cumulants do not.

#### 4.4.3 Walsh coefficients and background-averaged epistasis

The multivariate Bernoulli distribution also provides a probabilistic interpretation of the *Walsh coefficients* that are used to measure “background-averaged” epistasis [50, 41, 49, 60, 69]. The Walsh coefficients 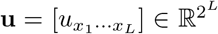, i.e. a vector indexed by binary strings, are defined by

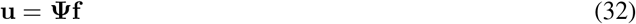

where 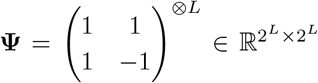 is a *Hadamard* matrix [50] and 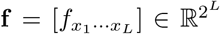 is the vector of fitness values indexed by binary strings. Equation (32) is known as the *Walsh-Hadamard* transformation, sometimes also called the *Walsh* or *Fourier-Walsh* transform; see [50, 41] for more details.

We prove that if the fitness values **f** are equal to probabilities **p** of a multivariate Bernoulli random variable (*X*_1_, …, *X*_*L*_), then the Walsh coefficients **u** are equal to the *moments* of (1 − 2*X*_1_, …, 1 − 2*X*_*L*_) ∈ {−1, 1}^*L*^, i.e. a linear transformation of the random variable (*X*_1_, …, *X*_*L*_) such that it takes values in {−1, 1}^*L*^ instead of {0, 1}^*L*^.

##### Theorem 2.

*Let* (*X*_1_, …, *X*_*L*_) ∈ {0, 1}^*L*^ *be distributed according to a multivariate Bernoulli distribution with general parameters* **f**, *and define Y*_*ℓ*_ = 1 − 2*X*_*ℓ*_ ∈ {−1, 1} *for ℓ* = 1, …, *L. Define* 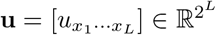 *as in* (32). *Then* 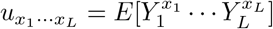.

To our knowledge, Theorem 2 gives the first probabilistic interpretation of the Walsh coefficients **u**. Interestingly, the Walsh coefficients **u** assume an *additive* fitness model [41, 50] while Theorem 2 requires that the fitness values **f** are equal to the probabilities **p**, an assumption corresponding to the *multiplicative* fitness model (Table 1).

Theorem 2 also provides a connection between the multivariate Bernoulli distribution and the circuit formulae used to quantify the geometry of a fitness landscape [59, 29, 60, 69], as the circuit formulae for a full genotype space are linear combinations of the Walsh coefficients; see [29] for details.

#### 4.4.4 Relationship to theoretical genetics models

We note that some previous works in theoretical genetics by Barton and Turelli (e.g. [136, 137, 138]) also model the genotype with a MVB. However, their approach is substantially different from ours. Barton and Turelli model linkage disequilibrium between *k* loci 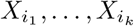 using the *k*-way central moment 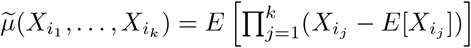 of the genotype distribution *P* (*X*_1_, …, *X*_*n*_). Barton and Turelli model epistasis using coefficients that are not related to the genotype distribution *P* (*X*_1_, …, *X*_*n*_). In contrast, we model epistasis with the natural parameters ***β*** of the genotype distribution *P* (*X*_1_, …, *X*_*n*_), as described in Section 4.4.1.

Interestingly, Barton and Turelli’s 3-way linkage disequilibrium term, i.e. 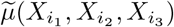, is equal to the 3-way 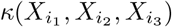 implicitly used by Kuzmin et al. [2, 42] to measure 3-way epistasis (see Section 4.4.2). This equivalence is because the *k*-way central moment 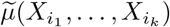 is equal to the *k*-way joint cumulant 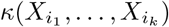 for *k* = 1, 2, 3. However, for *k* ≥ 4, the *k*-way linkage disequilibrium term used by Barton and Turelli is not equal to the *k*-way joint cumulant.

### 4.5 Simulating fitness values

We simulate fitness values *f*_x_ for genotypes **x** = (*x*_1_, …, *x*_*L*_) with *L* = 10 loci and *K*-way interactions using the following two different approaches. For both models, we divide all of the fitness values **f** by *f*_∅_ so that *f*_∅_ = 1.

#### Multiplicative fitness model

We draw interaction parameters *β*_*S*_ ∼ Uni(−0.5, 0.5) for each subset *S* ⊆ {1, …, *L*} of loci with size |*S*| ≤ *K*. We set the fitness *f*_x_ of genotype **x** = (*x*_1_, …, *x*_*L*_) as

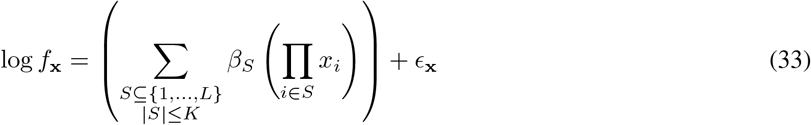

where *ϵ*_x_ ∼ *N* (0, *σ*^2^) are independent and identically distributed Gaussian random variables with mean zero and variance *σ*^2^.

#### NK model

We simulate fitness values **f** according to the NK model with the code used by [49]. Because [49] uses an additive fitness model, we exponentiate the fitness values from the NK model.

### 4.6 Epistasis between protein mutations

The analysis in Section 2.7 was performed using publicly available DMS data for the following proteins/RNA molecules:

- the *E*.*coli* metabolic protein folA [113];
- the Streptococcus pyogenes Cas9 (SpCas9) nuclease [114];
- the immunoglobulin-binding protein G domain B1 (GB1), expressed in Streptococcal bacteria [139, 140];
- the Omicron BA.1 variant of the SARS-CoV-2 virus, where fitness is measured relative to the Wuhan Hu-1 strain [141];
- the Entacmaea quadricolor fluorescent protein eqFP611, where fitness is measured in terms of fluorescence [142];
- the *Aequorea victoria* green fluorescent protein avGFP [143];
- the green fluorescent proteins (GFPs) from [144];
- yeast tRNA [145]; and
- the *Chlamydomonas reinhardtii* flavin mononucleotide (FMN)-based fluorescent protein CreiLOV [146].

We note that the fitness landscape of each protein may be measured using different scales. To make it possible to compare the fitness landscape of each protein, we make the following transformations to the observed fitness values **f**. For proteins whose fitness values are measured multiplicatively (resp. additively), we divide (resp. subtract) all fitness values *f* by the wild-type fitness value *f*_∅_. Moreover, for proteins whose fitness values are measured additively, we then exponentiate the fitness values (i.e. *f* → exp^*f*^) to convert the fitness values to a multiplicative scale. This allows us to compare the multiplicative epistasis measure *ϵ*^*M*^ with the chimeric epistasis measure *ϵ*^*C*^, which implicitly assumes fitness values are measured using a multiplicative scale.

For each protein and each interaction order *K*, we compute the multiplicative (resp. chimeric) measure *ϵ*^*M*^ (resp. *ϵ*^*C*^) across all *K*-tuples of mutational events. We note that for some proteins, the fitness of multiple mutations at a given locus is measured (e.g. all 3 possible base pair substitutions at a locus, or all 19 possible amino acid substitutions); for these proteins, we consider each possible mutation at a given genetic locus as a separate mutational event. Furthermore, following [139], we only compute the epistasis measure for a given *K*-tuple of mutational events if all fitness values for the 2^*K*^ genotypes are greater than a threshold *ϵ*, which we set to *ϵ* = 0.01 as in [139].

**Figure S1:**
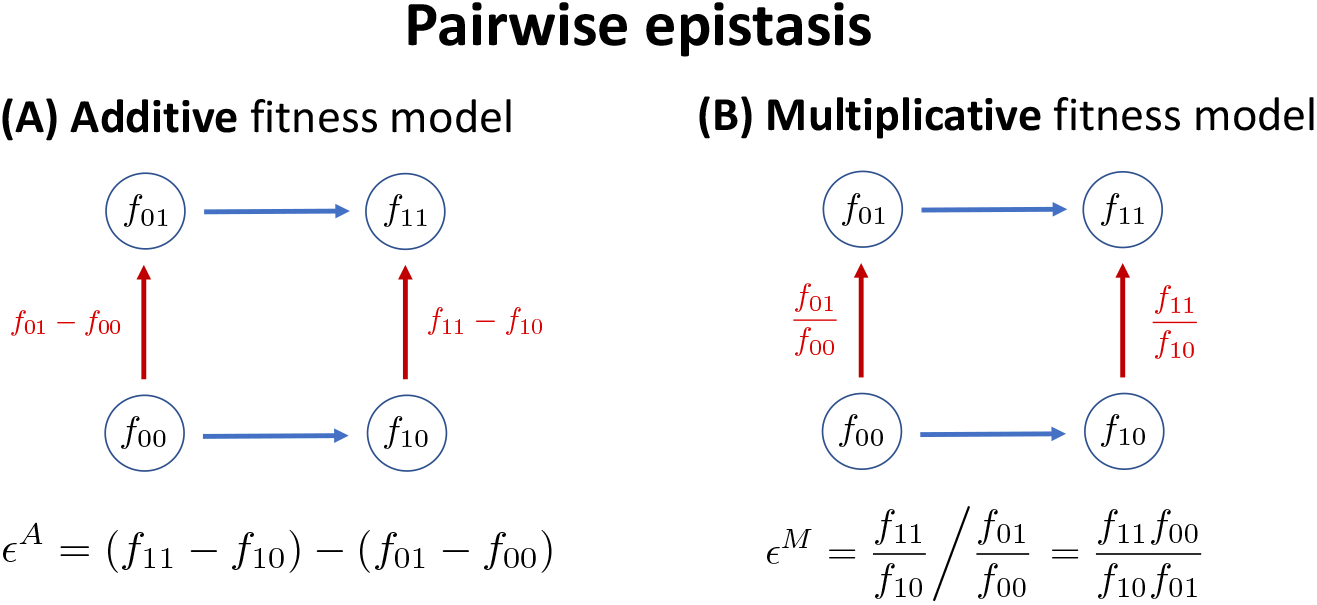
Pairwise epistasis is computed by the change in the fitness effect of a mutation in one locus across the two possible genetic backgrounds of the other locus (shown with the two red lines). The computation of the fitness effect and the change in the fitness effect depends on the fitness model. (**A**) In an additive fitness model, the mutational effect is computed with subtraction, while (**B**) in a multiplicative fitness model they are computed with a ratio.

## Supplementary Text

### A Comparing the chimeric and multiplicative formulae

For pairwise epistasis, the sign of the chimeric measure *ϵ*^*C*^ is always equal to the sign of the multiplicative epistasis measure *ϵ*^*M*^ (Proposition 1). Nevertheless, the chimeric epistasis measure may sometimes overstate or understate the degree of epistasis, as we demonstrate below.

**Example 1**. *Consider the two following scenarios under a multiplicative fitness model:*

1. *f*_11_ = 0.8, *f*_10_ = *f*_01_ = 1,*versus*
2. *f*_11_ = 0.2, *f*_10_ = *f*_01_ = 0.5.

*In both scenarios, the multiplicative epistasis measure is given by ϵ*^*M*^ = 0.8 *<* 1, *indicating the same degree of negative epistasis. However, the chimeric epistasis measure ϵ*^*C*^ *gives a different conclusion, namely that the first scenario (ϵ*^*C*^ = − 0.2 *<* 0*) has a larger degree of negative epistasis compared to the second scenario (ϵ*^*C*^ = − 0.05 *<* 0).

For higher-order epistasis, the sign of the chimeric epistasis measure is not guaranteed to be equal to the sign of the multiplicative epistasis measure, as we demonstrate below.

**Example 2**. *Consider a genotype with L* = 3 *loci and the following fitness values under the multiplicative model:*

1. *f*_1_ = 0.5, *f*_2_ = 1.0, *f*_3_ = 0.25,
2. *f*_12_ = 0.5, *f*_13_ = 1, *f*_23_ = 0.1,
3. *f*_123_ = 0.4

*Then there is no three-way epistasis using the multiplicative epistasis formula, i*.*e*.,

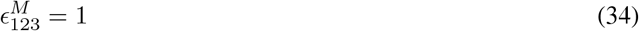

*However the chimeric three-way epistasis measure* 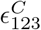 *is given by*

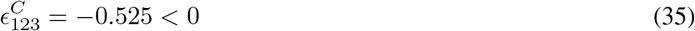

*and would indicate a negative three-way interaction*.

*If instead we have f*_1_ = 0.1 *and f*_123_ = 2 *then there is still no three-way epistasis according to the multiplicative measure, i*.*e*. 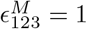, *but the chimeric three-way measure would incorrectly indicate a* positive *three-way interaction with* 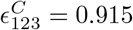.

### B Proof of Proposition 1

#### Proposition 1.

*Let f*_00_, *f*_01_, *f*_10_, *f*_11_ ∈ ℝ *be real numbers. Let* 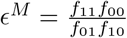 *and* 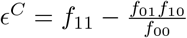. *Then*

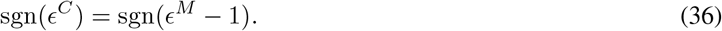

*Proof*. We have that

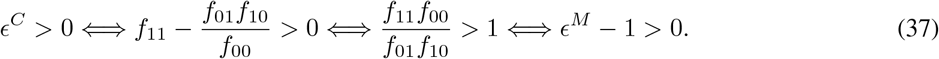

Thus, *ϵ*^*C*^ *>* 0 if and only if *ϵ*^*M*^ − 1 *>* 0. By similar logic, we have that *ϵ*^*C*^ = 0 (resp. *ϵ*^*C*^ *<* 0) if and only if *ϵ*^*M*^ − 1 *>* 0 (resp. *ϵ*^*M*^ − 1 *<* 0). It follows that sgn(*ϵ*^*C*^) = sgn(*ϵ*^*M*^ − 1).

### C Proof of Theorem 1

#### Theorem 1.

*Let* (*X*_1_, …, *X*_*L*_) ∈ {0, 1}^*L*^ *follow the multivariate Bernoulli distribution with general parameters* **p** *and natural parameters* ***β***. *Let* 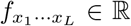 *be real numbers for each* (*x*_1_, …, *x*_*L*_) ∈ {0, 1} *such that* 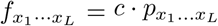 *for some constant c* ∈ ℝ. *Then we have*

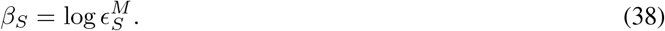

*Proof*. We proceed by induction on the size |*S*| of *S*. For our base case we assume that |*S*| = 2. Writing *S* = {*i*, }*j*, then we have that

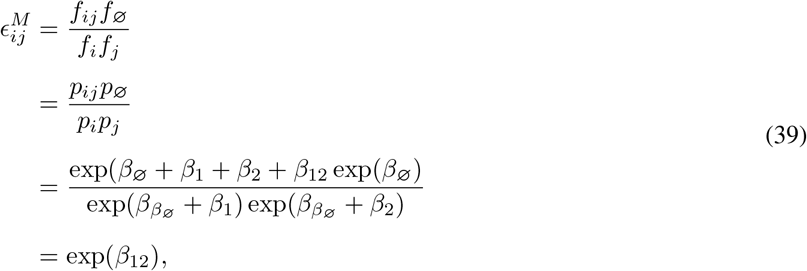

where in the second equality we use that the fitness values *f* are proportional to the probabilities *p* of the multivariate Bernoulli, and in the third equality we use the definition (23) of the natural parameters ***β***.

Next for the inductive hypothesis we assume that 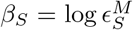 holds for all |*S*| *< K*. We will prove the equation holds for |*S*| = *K*. Without loss of generality assume that *S* = {1, …, *K*}. Then from (18) we have

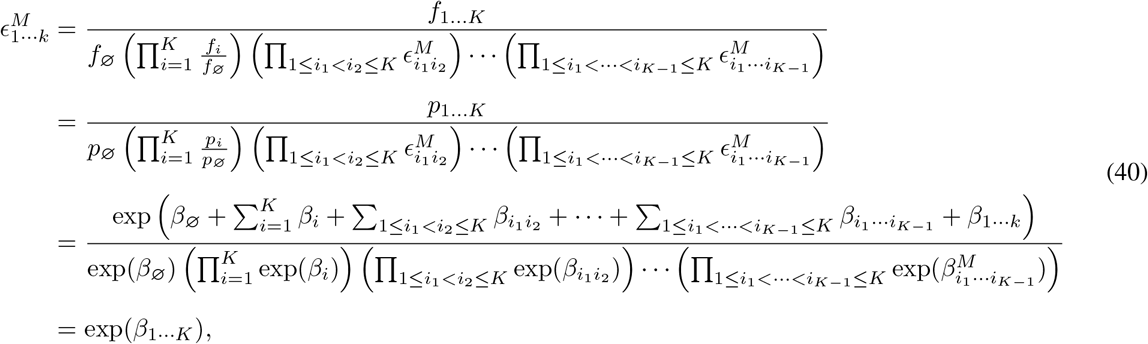

which completes the proof.

### D Transformation between parameterizations of the multivariate Bernoulli

Let (*X*_1_, …, *X*_*n*_) ∈ {0, 1}^*n*^ be distributed according to a multivariate Bernoulli distribution with natural parameters ***β***, mean parameters ***µ***, and general parameters **p**. [50, 82] give two formulae relating these different parametrizations. These equations are defined in terms of the *Kronecker product A* ⊗ *B* of matrices *A, B*. For shorthand, we write *A*^⊗*n*^ = *A* ⊗ *A* … ⊗ *A* as the Kronecker product of a matrix *A* with itself *n* times. Note that if *A* has size *s × t* then *A*^⊗*n*^ has size *s*^*n*^ *× t*^*n*^.

First, Equation (13) of [50] gives the following formula relating the natural parameters ***β*** and the general parame-ters **p**:

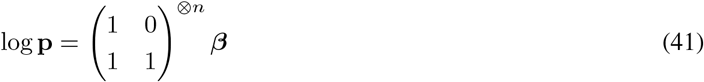

where log is taken entry-wise. Note that 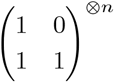 is a square matrix of size 2^*n*^ *×* 2^*n*^.

Second, [82] gives the following formula relating the general parameters **p** and the mean parameters ***µ***:

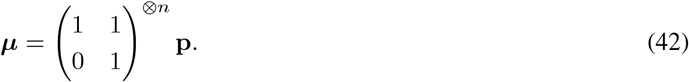

### E Proof of Theorem 2

We require the following lemma.

#### Lemma 1.

*Let* (*X*_1_, …, *X*_*L*_) ∈ {0, 1}^*L*^ *be distributed according to a multivariate Bernoulli distribution with moments* ***µ***^*C*^. *Define Y*_*ℓ*_ = 1 − 2*X*_*ℓ*_ ∈ {−1, 1} *for ℓ* = 1, …, *L. Let* 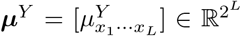 *be a vector with entries*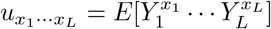. *Then we have*

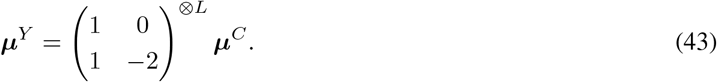

*Proof*. We proceed by induction on *L*. For ease of notation, we define 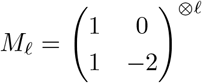. The base case, *L* = 1, is equivalent to

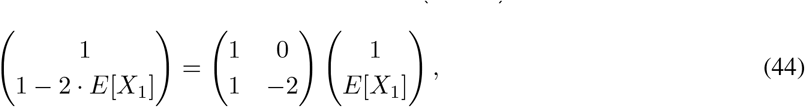

which holds by inspection.

Now for the inductive step, we assume (43) holds for *L* − 1 and we will show (43) holds for *L*. Define *A* = {(*a*_1_,*· · ·, a*_*L*_ ∈ {0, 1}^*L*^ : *a*_1_ = 0} and *B* = {(*a*_1_,*· · ·, a*_*L*_ ∈ {0, 1}^*L*^ : *a*_1_ = 1}. Then the first 2^*L*−1^ entries of ***µ***^*Y*^, ***µ***^*C*^ are indexed by *A*, and the second 2^*L*−1^ entries are indexed by *B*. Thus, define 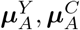 as the first 2^*L*−1^ entries of ***µ***^*Y*^, ***µ***^*C*^, respectively, and define 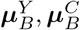 similarly.

For (0, *a*_2_,…, *a*_*L*_) ∈ *A*, we have that

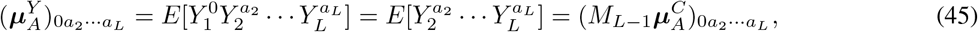

where in the last equality we use the inductive hypothesis on the *L*− 1 random variables (*X*_2_,…, *X*_*L*_). Similarly for (1, *a*_2_,…, *a*_*L*_) ∈ *B*, we have

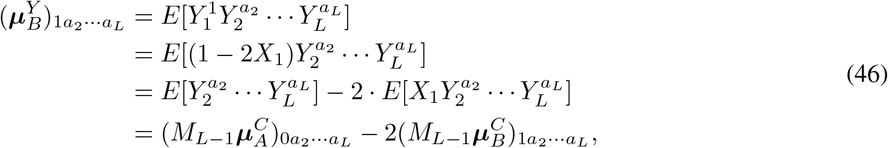

where again in the last equality we use the inductive hypothesis. Writing these two equalities in matrix form:

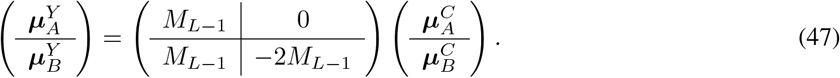

By definition of the Kronecker product, the matrix in (47) is equal to

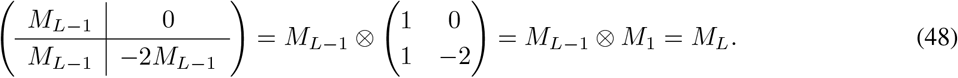

Thus, ***µ***^*Y*^ = *M*_*L*_***µ***^*C*^, completing the proof.

#### Theorem 2.

*Let* (*X*_1_, …, *X*_*L*_) ∈ {0, 1}^*L*^ *be distributed according to a multivariate Bernoulli distribution with general parameters* **f**, *and define Y*_*ℓ*_ = 1 − 2*X*_*ℓ*_ ∈ {−1, 1}. *Define* 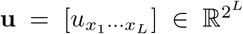 *as in* (32). *Then*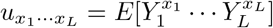.

*Proof*. Let ***µ***^*C*^ be the mean parameters of *X*. We define the vector 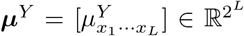 with entries 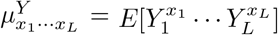.By Lemma 1 and equation (42), we have

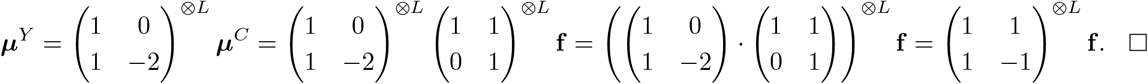

### F Relationship between MVB and case-control GWAS

Suppose we are given genotype (*X*_1_, *X*_2_) ∈ {0, 1}^2^ and (binary) disease status *D* ∈ {0, 1}. Then the joint random variable (*X*_1_, *X*_2_, *D*) follows a MVB distribution, where the log-probability log *P* (*X*_1_, *X*_2_, *D*) is given by the following expression in terms of the natural parameters ***β***:

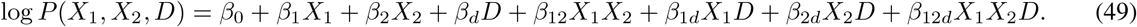

#### Logistic regression

In the logistic regression approach for measuring pairwise interactions, one fits a model of the form

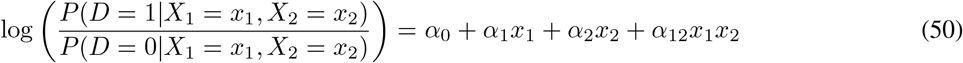

and measures pairwise interactions with the interaction term *α*_12_.

To relate the MVB and logistic regression, we rewrite the LHS of 50 as log *P* (*X*_1_ = *x*_1_, *X*_2_ = *x*_2_, *D* = 1)) − log *P* (*X*_1_ = *x*_1_, *X*_2_ = *x*_2_, *D* = 0) and plug in 49:

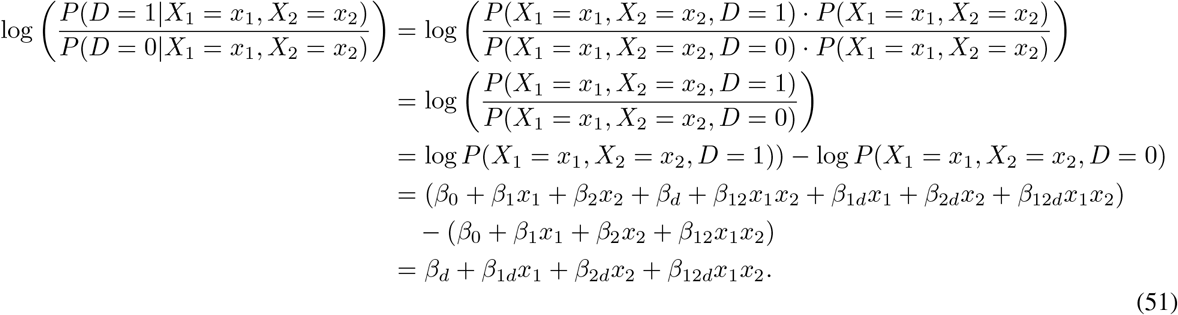

By equating coefficients with (50), it follows that *α*_12_ = *β*_12*d*_. That is, the 3-way interaction term *β*_12*d*_ in the MVB is equal to the logistic regression interaction term *α*_12_.

#### Conditional independence testing

We start by describing the conditional independence test. Let 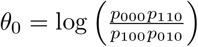 be the log-odds ratio of *X*_1_ and *X*_2_ conditioned on *D* = 0, where we use 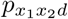 as shorthand for *P* (*X* = *x*_1_, *X*_2_ = *x*_2_, *D* = *d*). Note that *θ*_0_ = 0 if and only if *X*_1_ and *X*_2_ are independent conditioned on *D* = 0. Similarly, we define 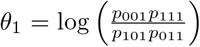 as the log-odds ratio of *X*_1_ and *X*_2_ conditioned on *D* = 1, so that *θ*_1_ = 0 if and only if *X*_1_ and *X*_2_ are independent conditioned on *D* = 1.

The conditional independence test is testing the null hypothesis *H*_0_ : *θ*_0_ = *θ*_1_ = 0. To relate the conditional independence test to the MVB, we plug (49) into the formula for *θ*_0_ and simplify, which yields:

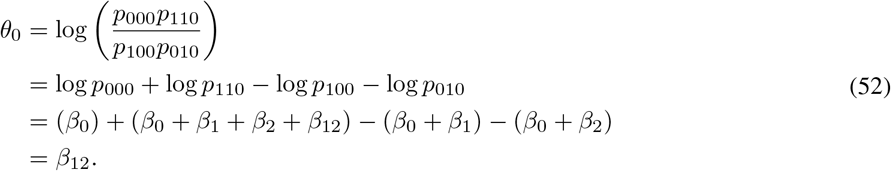

A similar computation for the formula for *θ*_1_ yields

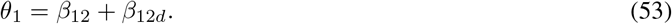

Thus, the conditional independence null hypothesis *H*_0_ : *θ*_0_ = *θ*_1_ = 0 is equivalent to the null hypothesis

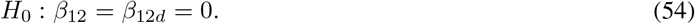

In other words, the conditional independence test is testing whether the 2-way interaction *β*_12_ (i.e. the *marginal* interaction) and 3-way interaction term *β*_12*d*_ are equal to 0.

#### Computing interaction terms *β* from GWAS data

From Theorem 1, each MVB interaction term is equal to the corresponding log-multiplicative measure, i.e. *β* = log *ϵ*^*M*^, where the fitness values 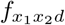 in the definition of the epistasis measure *ϵ*^*M*^ are proportional to the genotype probability 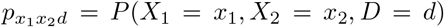, i.e. 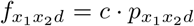 for some *c >* 0. Given observational GWAS data, the genotype probability 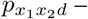 and thus the fitness values – are estimated by the empirical frequency of genotypes in the data with loci *X*_1_ = *x*_1_, *X*_2_ = *x*_2_ and disease status *D* = *d*.

### G Pairwise and higher-order epistasis measure comparison

The pairwise chimeric measure 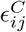 approximates the pairwise multiplicative epistasis measure 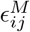 under certain con-ditions. Specifically, if the double mutant fitness *f*_*ij*_ and the product *f*_*i*_*f*_*j*_ of the single-mutant fitness values are both close to 1, i.e. *f*_*ij*_ ≈ 1 and *f*_*i*_*f*_*j*_ ≈ 1, then the pairwise chimeric epistasis measure 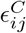 is approximately equal to the pairwise log-multiplicative measure log 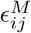. To see this, note that

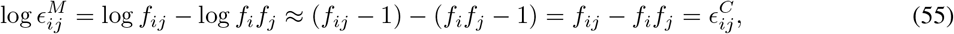

where we use the approximation that log *c* ≈ *c* − 1 if *c* ≈ 1.

We empirically assessed (Figure S2) how closely the the pairwise chimeric epistasis measure 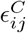 approximates the interaction parameter *β*_*ij*_ (which is equal to the pairwise log-multiplicative measure log 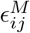), using the simulated fitness values **f** from the multiplicative fitness model (Section 2.4, pairwise interactions *K* = 2 with noise parameter *σ* = 0). We observe that the chimeric measure has small error 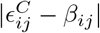 when both the double mutant fitness *f*_*ij*_ and product *f*_*i*_*f*_*j*_ of single mutant fitness values are close to 1. However, the error gets much larger when either *f*_*ij*_ or *f*_*i*_*f*_*j*_ are not close to 1, which agrees with when the the approximation in (55) is valid. Moreover, the multiplicative and chimeric measures diverge substantially for higher-order interactions, with the correlation between the two measures approaching zero as the interaction order increases (Figure S3).

This analysis demonstrates that for pairwise interactions, the chimeric epistasis measure 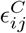 is an accurate measure of interactions in a multiplicative fitness model in certain settings.

**Figure S2:**
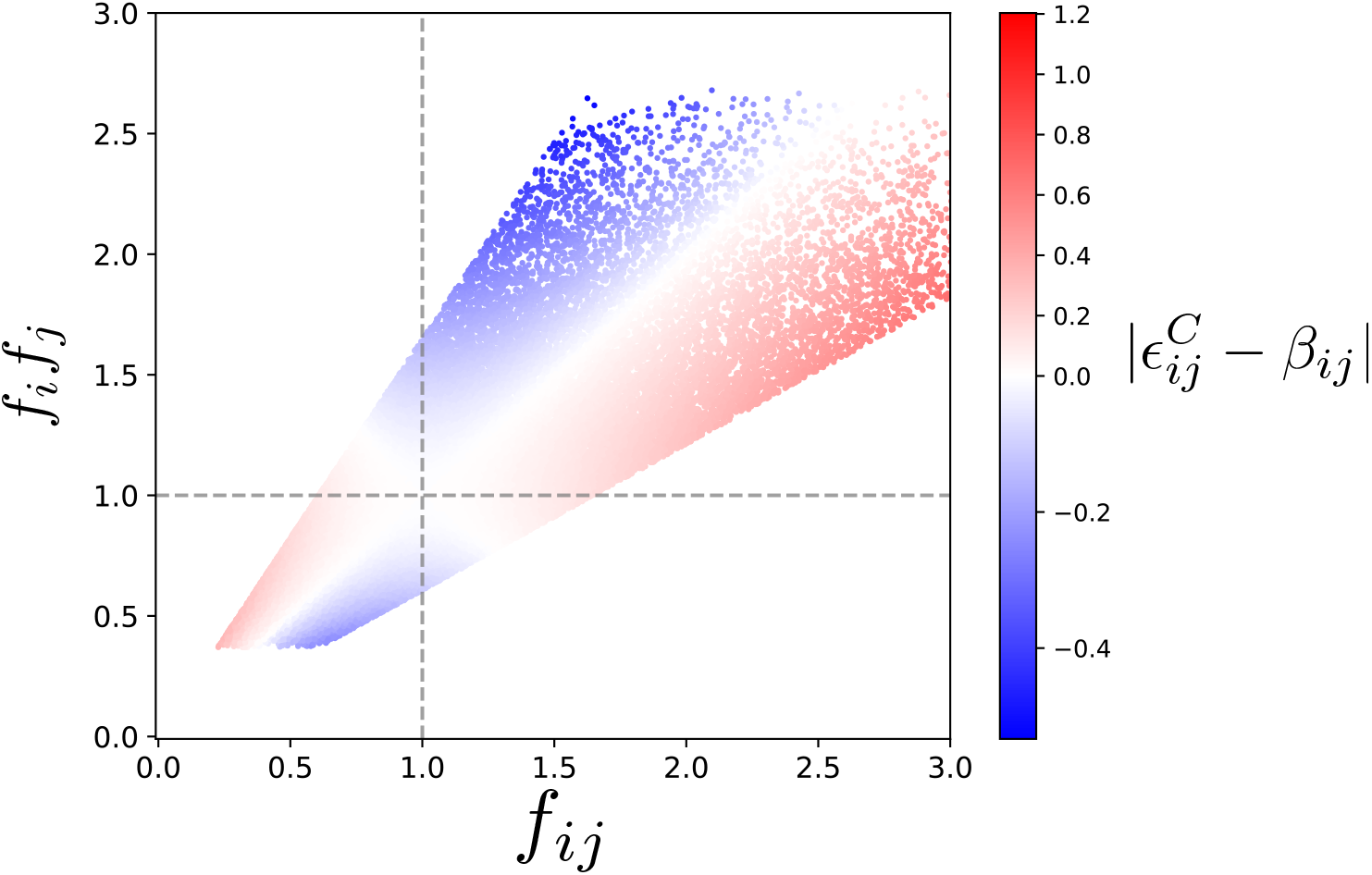
Double mutant fitness *f*_*ij*_ versus product *f*_*i*_*f*_*j*_ of single mutant fitness values for each pair (*i, j*) of loci across all simulated instances of the multiplicative fitness model in Section 2.4 (pairwise interactions with noise parameter *σ* = 0). Points are colored by the difference 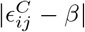 between the chimeric epistasis measure 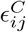 and true interaction parameter *β*.

### H Yeast reanalysis

#### H.1 Reproducing previous results

We gathered fitness data from the supplemental tables of [42] and [2] in a way that was consistent with how the data were described in the corresponding publications. To reproduce results from [42], we used data from Supplementary Tables S1, S3, and S5. To reproduce those from [2] we used Table S1 along with Data File S1 from [31] which contains three tables (SGA NxN.txt, SGA ExN.txt, SGA ExE.txt).

Following the formula and notation from the supplement of [2], we recalculated the trigenic interaction scores *τ*_*ijk*_ reported by [2, 42] (i.e. the chimeric three-way measures 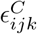) using the following formula:

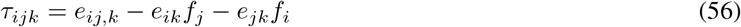

where

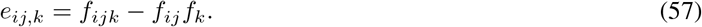

Here, *e*_*jk*_ and *e*_*ik*_ are the reported pairwise interactions between knockout mutant *j, k* and *i, k*, respectively. We obtained the values of *f*_*i*_, *f*_*j*_, *f*_*k*_, *f*_*ij*_, *f*_*ijk*_, *e*_*ik*_, and *e*_*jk*_ directly from the data tables of [2, 42]. We recomputed the trigenic interaction score *τ*_*ijk*_ for 74% and 81% of the triple knockout mutants (*i, j, k*) reported in [42] and [2], respectively, as some values were either missing or were “NaN”. Our recalculated trigenic interaction scores were highly similar to reported values: the distributions were nearly identical and values were highly correlated (Figure S4), with a Pearson correlation of 0.9974 and 0.9809 for the data from [42] and [2], respectively.

We note that plugging (57) into (56) yields the following expression for the trigenic interaction score *τ*_*ijk*_:

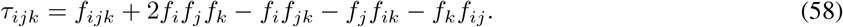

Computing the trigenic interaction score *τ*_*ijk*_ using (58) should be identical to computing it using (56) (Figure S4).

However, we find that the scores calculated using (58) are quite different compared to the scores reported by [42, 2] (Figure S5). This discrepancy is due to the fact that the reported values of *f*_*ik*_ and *f*_*jk*_ are not equal to *f*_*i*_*f*_*k*_ + *e*_*ik*_ and *f*_*j*_*f*_*k*_ + *e*_*jk*_, respectively. Thus, to make our reanalysis consistent with the reported trigenic interaction scores from [2, 42], we recompute the trigenic interaction score using (56).

We computed the three-way multiplicative measure 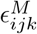 as

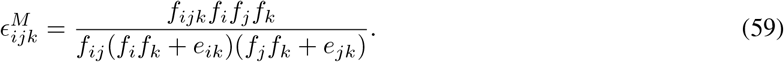

**Figure S3:**
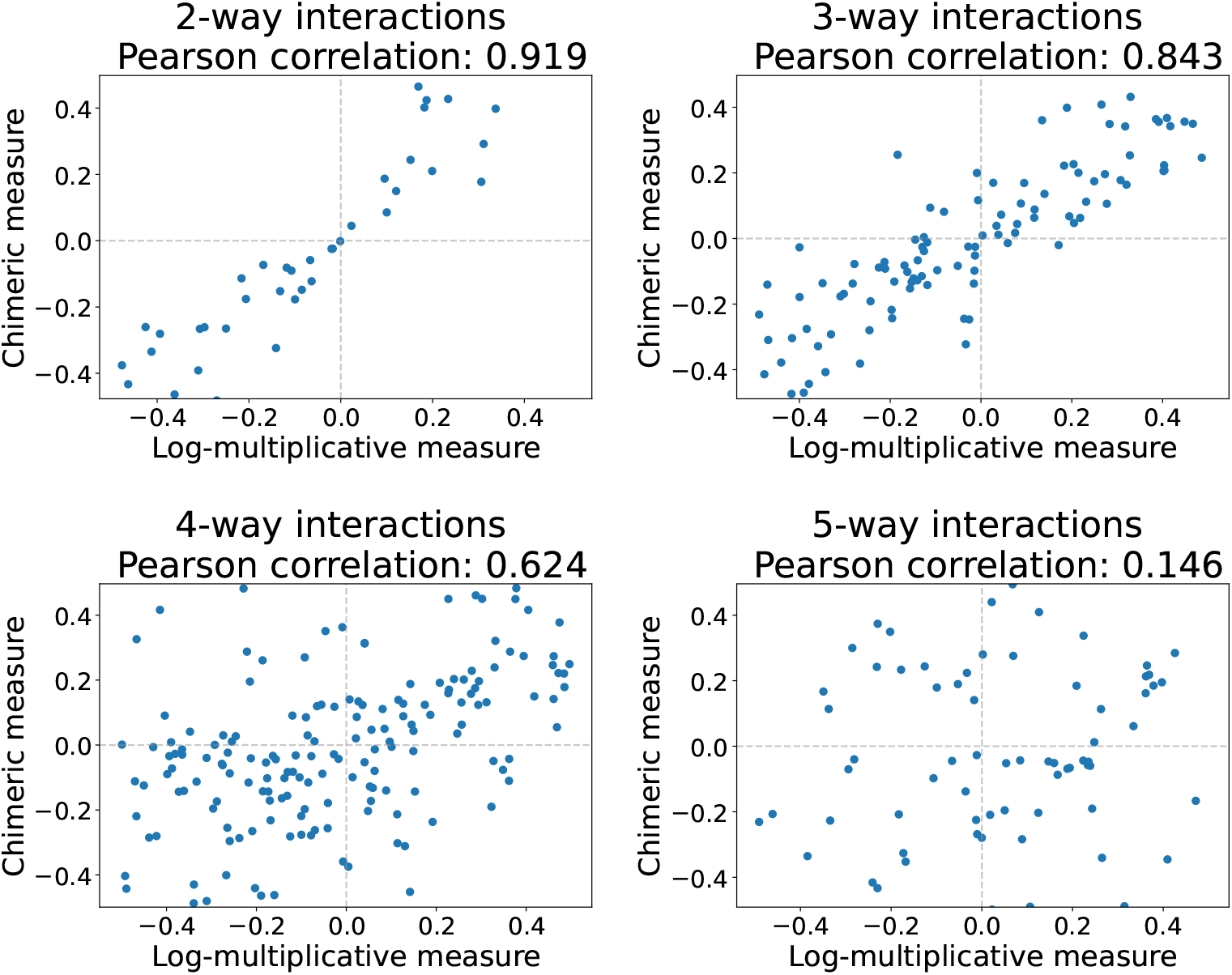
Pearson correlation *ρ*(*ϵ*^*C*^, log *ϵ*^*M*^) between chimeric measure *ϵ*^*C*^ and multiplicative measure *ϵ*^*M*^ for different interaction orders *K* and each *K*-tuple of loci across all simulated instances of the multiplicative fitness model in Section 2.4 (pairwise interactions with noise parameter *σ* = 0). The *x*- and *y*-axis ranges are [min *ϵ*^*M*^, max *ϵ*^*M*^ ].

In the denominator, we compute the double-mutant fitness values *f*_*ik*_ and *f*_*jk*_ indirectly due to the aforementioned issues with using the reported double-mutant fitness values *f*_*ik*_ and *f*_*jk*_. Our data handling procedures and analysis are located in our Github repository.

#### H.2 Enrichment analyses

##### Physical interactions

To test whether subsets of trigenic interactions were enriched for protein-protein interactions, we downloaded pairwise physical interaction data from BIOGRID (release 4.4.211) for Saccharomyces cerevisiae (strain S288c) and only considered physical interactions discovered using the following experimental system types: Affinity Capture-MS, Affinity Capture-Western, Two-hybrid, Reconstituted Complex, PCA, Co-purification, and Co-crystal Structure. Using this information, we classified a given set of three genes as having a shared protein-protein interaction if all three genes had a physical interaction with at least one other gene in the genome (not including the genes in the set of three).

**Figure S4:**
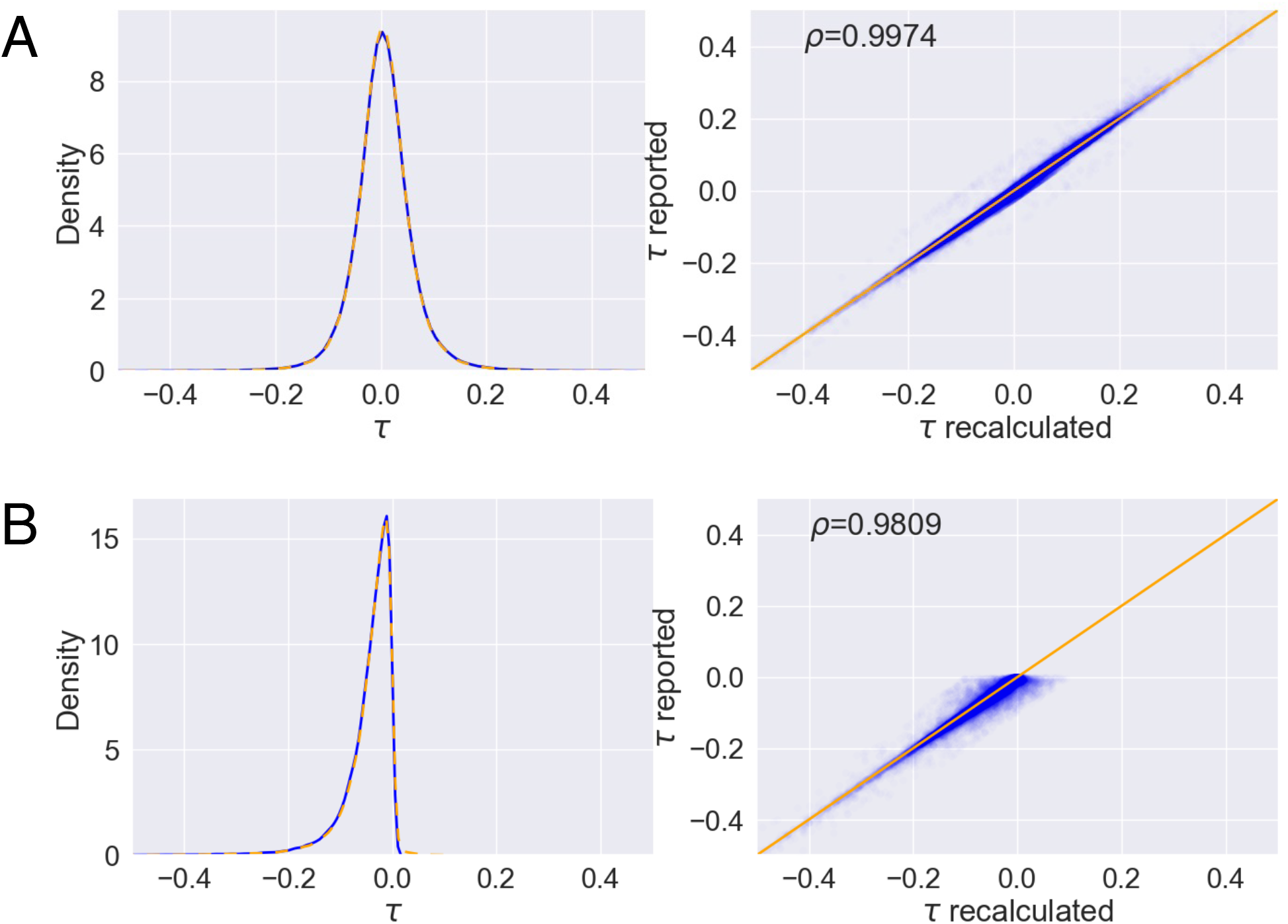
**(A)** Kuzmin et al 2020 [42]. Left: Density of the reported trigenic interaction score *τ*_*ijk*_ (blue) and our recalculation of *τ*_*ijk*_ using (56) from fitness data (orange). Right: Calculation of the Pearson correlation between the reported value of *τ*_*ijk*_ and our recalculation of *τ*_*ijk*_ using (56). **(B)** Same analysis as in (A) for data from Kuzmin et al 2018 [2]. Note that for this analysis, only negative values of *τ*_*ijk*_ were reported.

##### Coexpression

From COXPRESSdb [147], we downloaded union-type coexpression data, which integrates RNAseq and microarray coexpression data, for S. cerevisiae. From these data, we only considered a pair of genes as significantly coexpressed if they had a Z-score greater than or equal to 3. To test whether subsets of trigenic interactions were enriched for coexpressed genes, we classified a given set of three genes as coexpressed if at least two of three possible gene pairs within the set had a coexpression Z-score greater than or equal to three.

##### GO enrichment

We downloaded GO Slim mappings from the Saccharomyces Genome Database and only considered GO terms corresponding to biological processes (i.e. with a GO aspect of “P” in the GO Aspect column of go slim mapping.tab). We then classified a given set of three genes as sharing a GO term if all three genes had at least one GO term in common.

#### H.3 Trigenic interaction fraction

Kuzmin et al. [42] quantify functional redundancy between paralogs using a quantity that they call the *trigenic interaction fraction*. The trigenic interaction fraction is equal to the number of trigenic interactions involving both paralogs divided by the sum of numerator and the number of digenic interactions involving at least one of the paralogs. Kuzmin et al. [42] hypothesize that paralogs with higher trigenic interaction fractions are more redundant, whereas those with low trigenic fractions have undergone subfunctionalization.

We compared the trigenic interaction fraction computed using the chimeric formula to the trigenic interaction fraction computed using the multiplicative formula (Figure S10). We observe that for most gene pairs, the trigenic interaction fraction is larger when it is computed using the multiplicative formula versus when it is computed using the chimeric formula. Additionally, for the 15 paralogs that have many additional trigenic interactions found using the multiplicative formula (highlighted in Figures 4C, D), we also observe elevated trigenic interaction fractions (Figure S10). This observation further supports the conclusion that the multiplicative measure uncovers additional functional redundancies between these paralogs.

**Figure S5:**
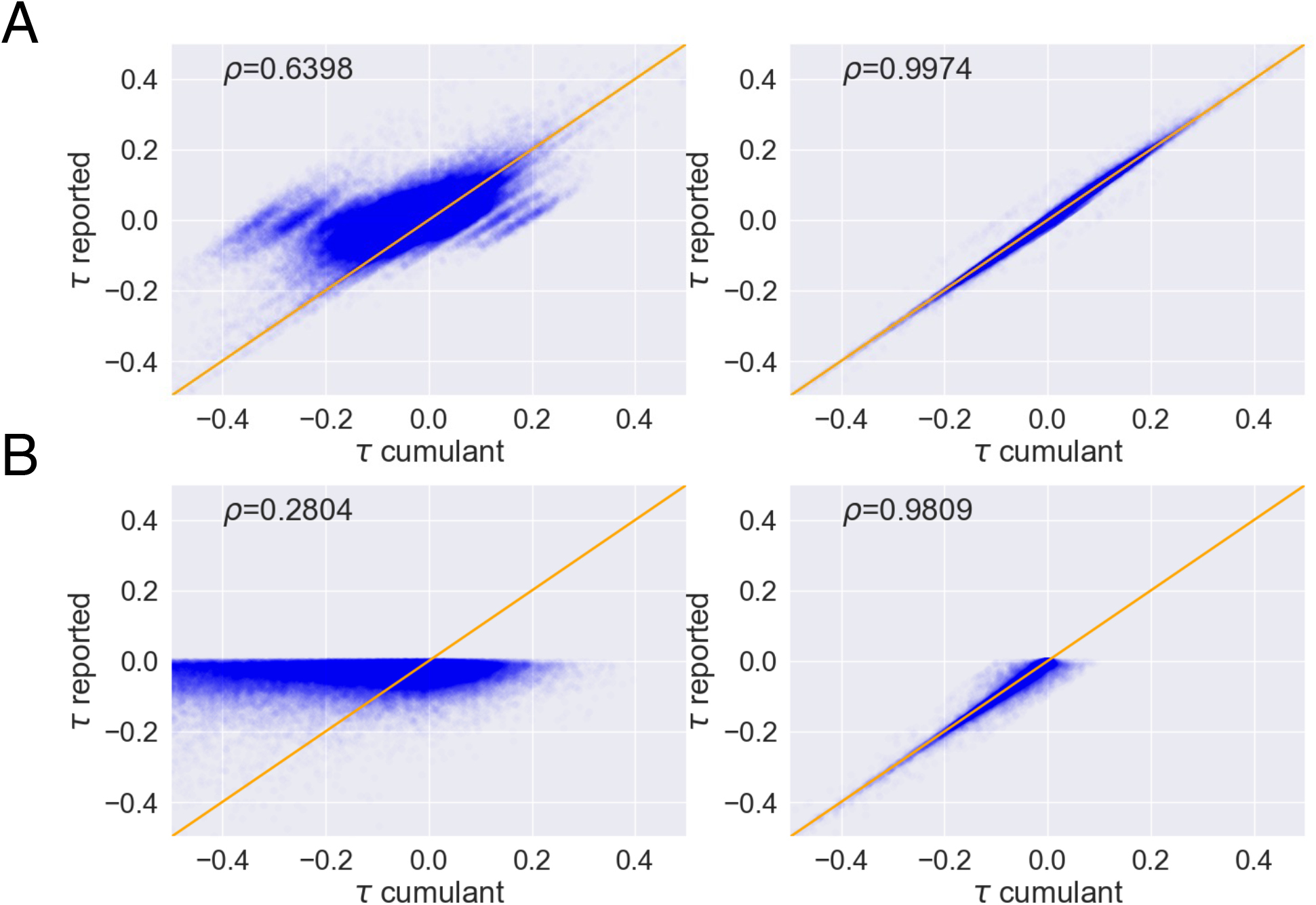
Same analysis as in Figure S4, except with the trigenic interaction score *τ*_*ijk*_ calculated using (58).

**Figure S6:**
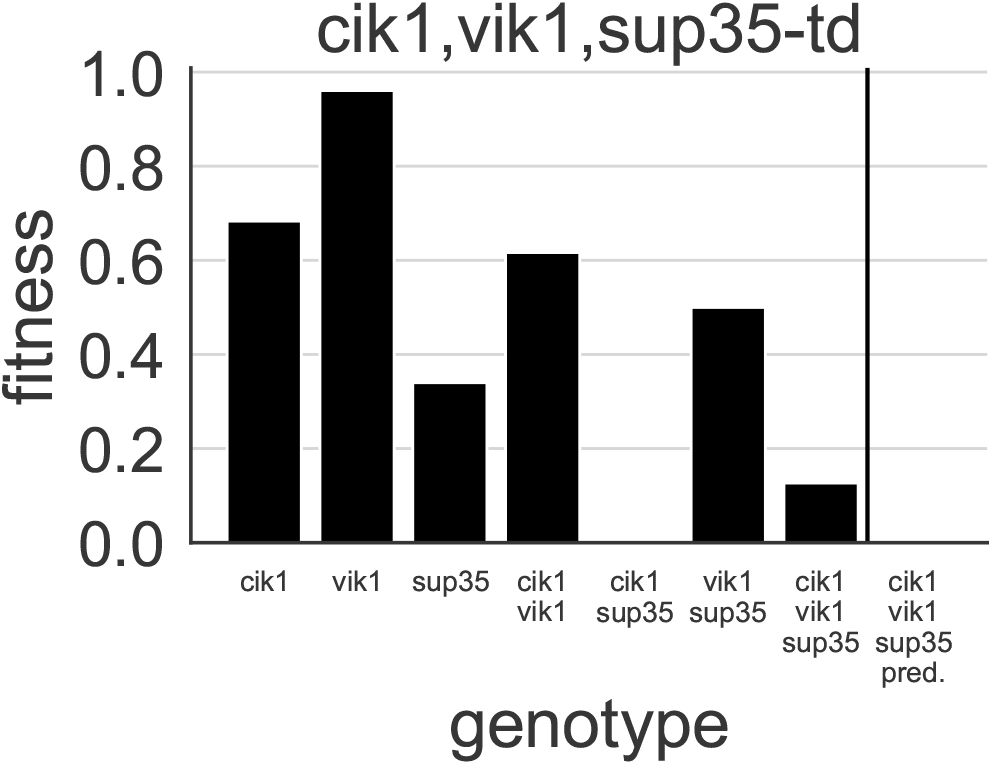
Single, double, and triple mutant fitnesses are shown for the cik1, vik1, sup35-td gene triple. On the right-hand side of the plot is the predicted triple-knockout fitness using all single mutant fitnesses and pairwise multiplicative epistasis measures 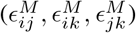.The observed triple-knockout fitness is substantially higher than predicted, indicating higher-order epistasis.

**Figure S7:**
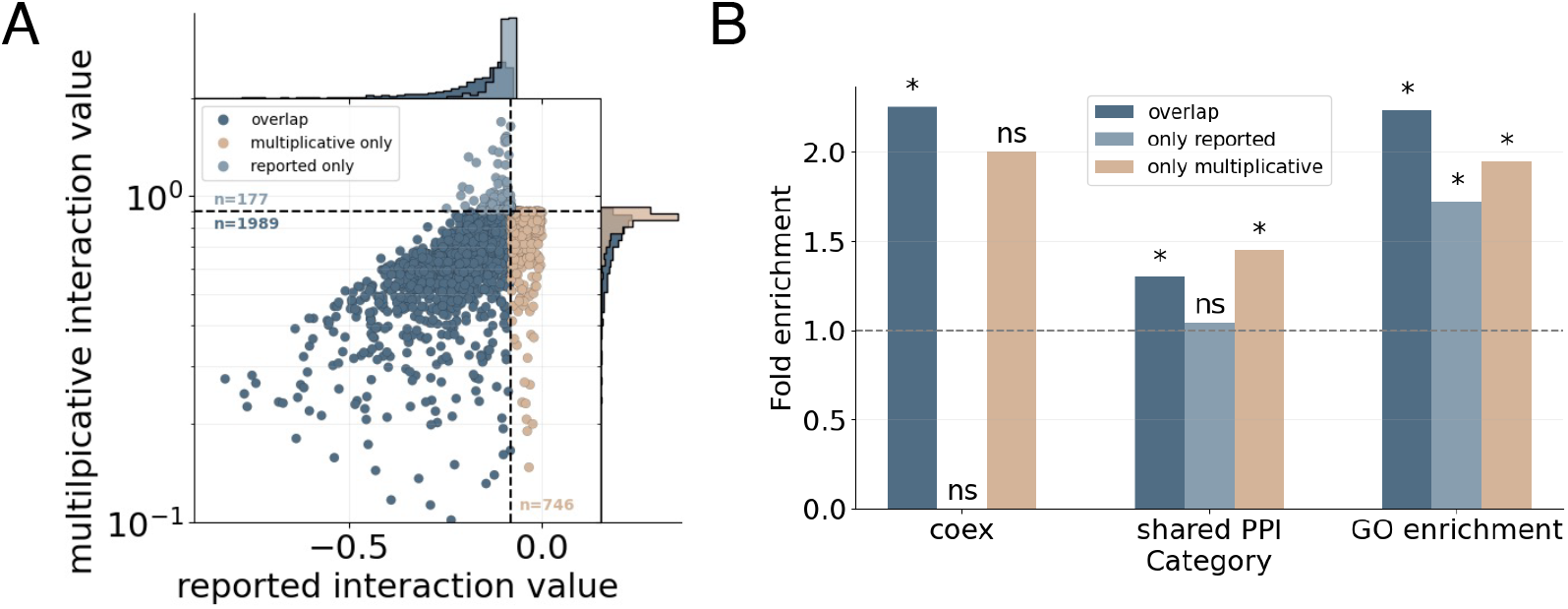
Comparison of negative trigenic interactions reported by Kuzmin et al 2018 [2] and those detected using the multiplicative epistasis formula. The analysis is identical to Figure 4 but using data from Kuzmin et al 2018 [2] instead of Kuzmin et al 2020 [42].

**Figure S8:**
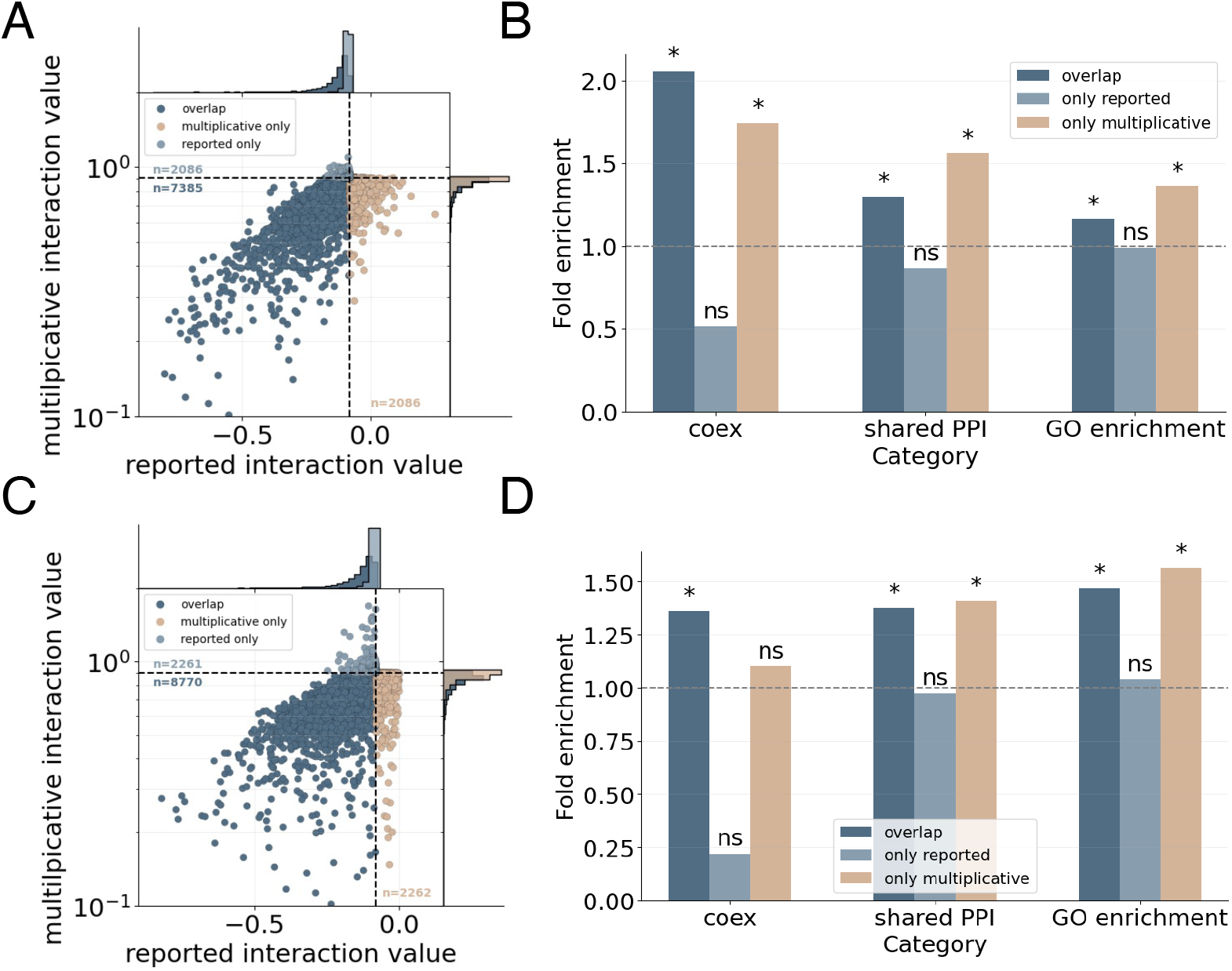
Comparison of negative trigenic interactions reported by [42] (top row) and [2] (bottom row) and those detected using the multiplicative formula. These analyses are identical to those reported in Figure 4 and Figure S7 except we do not use the reported *p*-value from [2, 42] to filter negative interactions (see Section 2.5).

**Figure S9:**
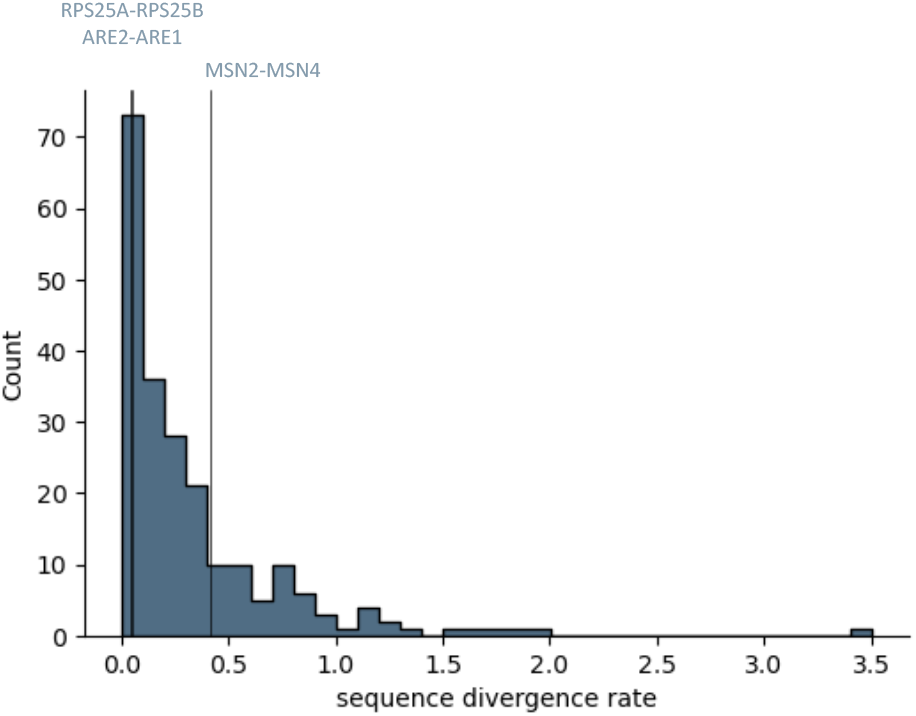
Distribution of sequence divergence rates between paralogs interrogated by [42]. RPS24A-RPS25B and ARE1-ARE2 have low sequence divergence rates of 0.041 and 0.051, respectively, whereas MSN2-MSN4 has a higher divergence rate of 0.41.

**Figure S10:**
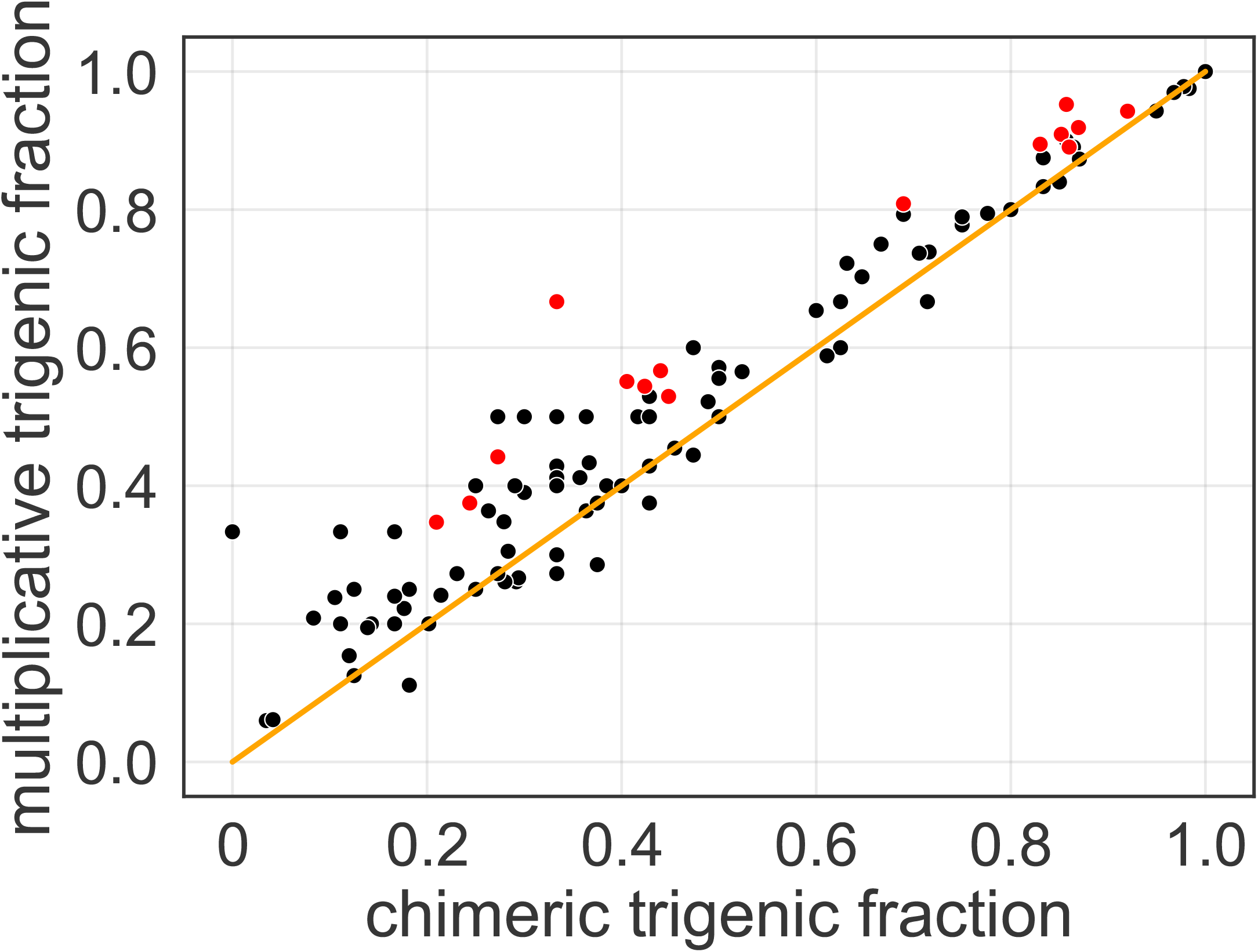
Comparison of trigenic fractions computed using the chimeric formula (x-axis) and the multiplicative formula (y-axis) for paralog pairs with at least 6 total interactions (trigenic and digenic). Red dots indicate the paralog pairs highlighted in Figure 4 which had many additional trigenic interactions according to the multiplicative measure. The orange line indicates the point at which trigenic fractions are equivalent across measures.

## Footnotes

1 While this parameter is typically called *N* (i.e. the “N” in “NK”), we use *L* to maintain consistency with the notation in this manuscript.

2 As this is sometimes a point of confusion: we note that triangles in a graph are sometimes referred to as higher-order structures [89]. However, as our simulation demonstrates, it is quite possible to have a triangle in a graph, i.e. three pairwise interactions, without having a genuine higher-order (3-way) interaction.

